# Self-Assembled Generation of Multi-zonal Liver Organoids from Human Pluripotent Stem Cells

**DOI:** 10.1101/2024.08.30.610426

**Authors:** Hasan Al Reza, Connie Santangelo, Abid Al Reza, Kentaro Iwasawa, Sachiko Sachiko, Kathryn Glaser, Alexander Bondoc, Jonathan Merola, Takanori Takebe

## Abstract

Distinct hepatocyte subpopulations are spatially segregated along the portal-central axis and critical to understanding metabolic homeostasis and liver injury. While several bioactive molecules have been described to play a role in directing zonal fates, including ascorbate and bilirubin, *in vitro* replication of zonal liver architecture has not been achieved to date. In order to evaluate hepatic zonal polarity, we developed a self-assembling zone-specific liver organoid culture by co-culturing ascorbate and bilirubin enriched hepatic progenitors derived from human induced pluripotent stem cells. We found that preconditioned hepatocyte-like cells exhibited zone-specific functions associated with urea cycle, glutathione synthesis and glutamate synthesis. Single nucleus RNA sequencing analysis of these zonally patterned organoids identifies hepatoblast differentiation trajectory that mimics periportal-, interzonal-, and pericentral human hepatocytes. Epigenetic and transcriptomic analysis showed that zonal identity is orchestrated by ascorbate or bilirubin dependent binding of histone acetyltransferase p300 (EP300) to methylcytosine dioxygenase TET1 or hypoxia-inducible factor 1-alpha (HIF1α). Transplantation of the self-assembled zonally patterned human organoids improved survival of immunodeficient rats who underwent bile duct ligation by ameliorating the hyperammonemia and hyperbilirubinemia. Overall, this multi-zonal organoid system serves as an *in vitro* human model to better recapitulate hepatic architecture relevant to liver development and disease.

---

The liver is a multi-faceted organ with a wide range of functions such as glycolysis and gluconeogenesis, lipogenesis and fatty acid oxidation, protein synthesis and xenobiotic catabolism. These divergent and complex functions are spatially segregated through compartmentalization into distinct regions called Zone 1, 2, 3 hepatocytes, based on their proximity from the central vein to the portal vein ^1,2^. For example, the maintenance of nitrogen level in the liver is precisely balanced between the input of nutrients and the output as ammonia, which is metabolized by the urea cycle, nitric oxide cycle, and glutamine synthesis ^3–5^. Urea cycle is primarily carried out in zone 1 and 2 hepatocytes, while glutamine synthesis takes place in zone 3 hepatocytes ^6^. On the other hand, the nitric oxide cycle is primarily maintained in zone 2 and 3 hepatocytes^7^. Citrulline is a key metabolic element necessary for these pathways and the levels of citrulline are maintained by the nitric oxide cycle, which is augmented by glutathione ^8^. As a consequence of this zonal segregation of hepatic functions, metabolic diseases tend to manifest within the particular zone in which the derangement is most significant.

While much of the expression patterns are conserved between rodent and human liver zonation, some variations in metabolic compartmentalization exists due to the differences in metabolic demands ^2^. Specifically, only 35.3% of all genes expressed in the human liver have similar expression to those in mice suggesting the existence of unique genetic networks and transcriptional regulation ^9,10^. One example is glutamine synthetase (GLUL) and carbamoylphosphate synthetase (CPS1) that participate in nitrogen metabolism, that overlap in pericentral and periportal regions, respectively in rat liver ^11^. However, the human liver contains an intermediate zone where neither enzyme is present. These inherent genetic and molecular differences necessitate a need for a human system to model development and disease that occur in patients.

Several studies indicate that zonation in the adult liver is maintained by the availability of nutrients and oxygenation status delimited by the portal and central veins. The candidate signaling cascade responsible for proper zonation of the pericentral hepatocytes is the Wnt family molecules, while the Hedgehog (Hh) and Notch signaling pathway are linked to periportal hepatocytes and cholangiocytes ^12^. Efforts to model this zonality are underway in animal and human cell culture with attempts to create oxygen ^13^, chemical ^14,15^, and genetic gradient conditions ^16^. However, the principal transcriptional regulators of zonal fate determination have yet to be defined, hindering the ability to effectively model zonal hepatocyte differentiation in human tissues.

Ascorbic acid, an essential antioxidant that is essential for hepatocyte development ^17^, regulates the expression of several Zone 1 specific liver genes. Periportal hepatocytes are principally responsible for functions including gluconeogenesis ^18^, cholesterol synthesis ^19^ and fatty acid oxidation ^20^ are potentiated by the activity of ascorbate, whereas lipogenesis, attributed to pericentral hepatocytes, is inhibited by ascorbic acid ^21,22^. Ascorbate is also known for the activation of *Tet1* (Tet methylcytosine dioxygenase 1) in the liver, which in turn activates Hh signaling essential for the activation of periportally-enriched gene expression in late embryogenesis ^12,20^. Collectively these lines of evidence support that stabilized ascorbic acid exposure can promote the expression of periportal specific metabolic pathway genes.

In contrast, bilirubin, a metabolic waste product made from heme ^23^, has a potential to enrich metabolic activities located in the pericentral areas. For example, bilirubin can promote the expression of Zone 3- enriched specific CYP enzymes directly ^24,25^ or indirectly through Wnt signaling activation^26,27^. Wnt activating role of bilirubin is explained by its pro-angiogenic properties thereby activating the Akt-NOS3 signaling pathway^28,29^. Moreover, bilirubin is known to activate both transcription and translation of HIF1α in even normoxic conditions to emulate the after-effects on exposure to hypoxic conditions^30^. Given that GLUL+ pericentral hepatocytes are HIF1α positive, these findings suggest that bilirubin sustains the expression of Zone 3 specific programs.

In order to employ differential inductive conditions, we developed dual liver tissue culture system by combining ascorbate- and bilirubin- enriched progenitors derived from human induced pluripotent stem cells (hiPSCs). Since the *L-gulonolactone oxidase (GULO)* gene lost the ability to synthesize ascorbate in humans due to pseudogenization ^31^, we took advantage of a tetracycline (TET) inducible active *Gulo* knock-in hiPSC ^32^, which was used to evoke periportal identity, whereas standard hiPSC lines were exposed to bilirubin to prime into pericentral lineage. The transcriptomic, epigenetic and functional profile of the generated organoids support the emergence of multizonal phenotypes and functions that ameliorate fatal total liver dysfunction upon transplantation in an immunosuppressed rodent model.

## Results

### Intracellular ascorbate specifies CPS1+ hepatocytes in Z1-HLOs

The Osteogenic Disorder Shionogi (ODS) rat is an animal models that has a genetic defect in hepatic L-gulonolactone oxidase with impaired hepatic metabolism ^33^. To test the importance of gulo in zonal identity, we first stained key periportal and pericentral markers, Glutaminase2 (GLS2) and Glutamate Synthetase (GS) with or without dietary ascorbic acid supplementation. 7∼14 days after ascorbic acid deprivation, GLS2+ve periportal hepatocyte fractions significantly reduced, whereas GS+ve pericentral fraction modestly increased (**Extended Data Fig. 1a**). To examine the role of gulo in zone-priming effect, we next employed stem cell differentiation models. We first established an inducible human iPSC cell line with the murine *Gulo* (*mGulo*) transgene and a P2A mCherry inserted into the AAVS1 locus with a tetracycline (TET)-inducible system^32^. These *mGulo* iPSCs were used to induce posterior foregut cells based on previously published protocols ^34–36^, and eventually generate putative Zone 1-like human liver organoids (Z1-HLOs) by continued Doxycycline (dox) induction beginning on Day 17 (Fig. 1a and Extended Data Fig. 1b). Presumptive mGulo expression was paralleled by the mCherry expression. In ascorbate deprived condition, iPSCs lacking the mGulo transgene failed to differentiate into HLO properly, whereas *mGulo* containing iPSCs generated healthy viable HLOs without ascorbate (Fig. 1b and Extended Data Fig. 1b). The expression of dox induced *mGulo* contributed to higher cellular antioxidant concentration and lower ROS levels when compared to exogenous ascorbate supplements in medium (Extended Data Fig. 1c, d, and e). These Z1-HLOs had higher expression of zone 1 liver genes such as *FAH (Fumarylacetoacetase), HPD (4-Hydroxyphenylpyruvate dioxygenase),* and *SCD (Stearoyl-CoA desaturase)* (Extended Data Fig. 1f). Compared to mGulo absent HLOs, the Z1-HLOs showed higher expression of urea cycle genes such as *ACSS2 (Acyl-coenzyme A synthetase 2), ASL (Argininosuccinate lyase), CPS1*, and *OTC (Ornithine transcarbamylase)* approaching to those in primary hepatocytes (PHH) (Fig. 1b). ELISA assay showed that the Z1-HLOs synthesized much higher levels of albumin compared to control HLOs and primary human hepatocytes (PHH), a characteristic prominent in periportal hepatocytes (**Extended Data Fig. 1g**). More importantly, the Z1-HLOs expressed CPS1 and ACSS2 proteins as verified by immunofluorescence and compared to control HLOs (**Extended Data Fig. 1h**). Overall, these data suggests that functional mGulo induction with elevated intracellular ascorbate primes the differentiation into CPS1+ periportal hepatocyte in Z1-HLOs.

**Fig.1.**
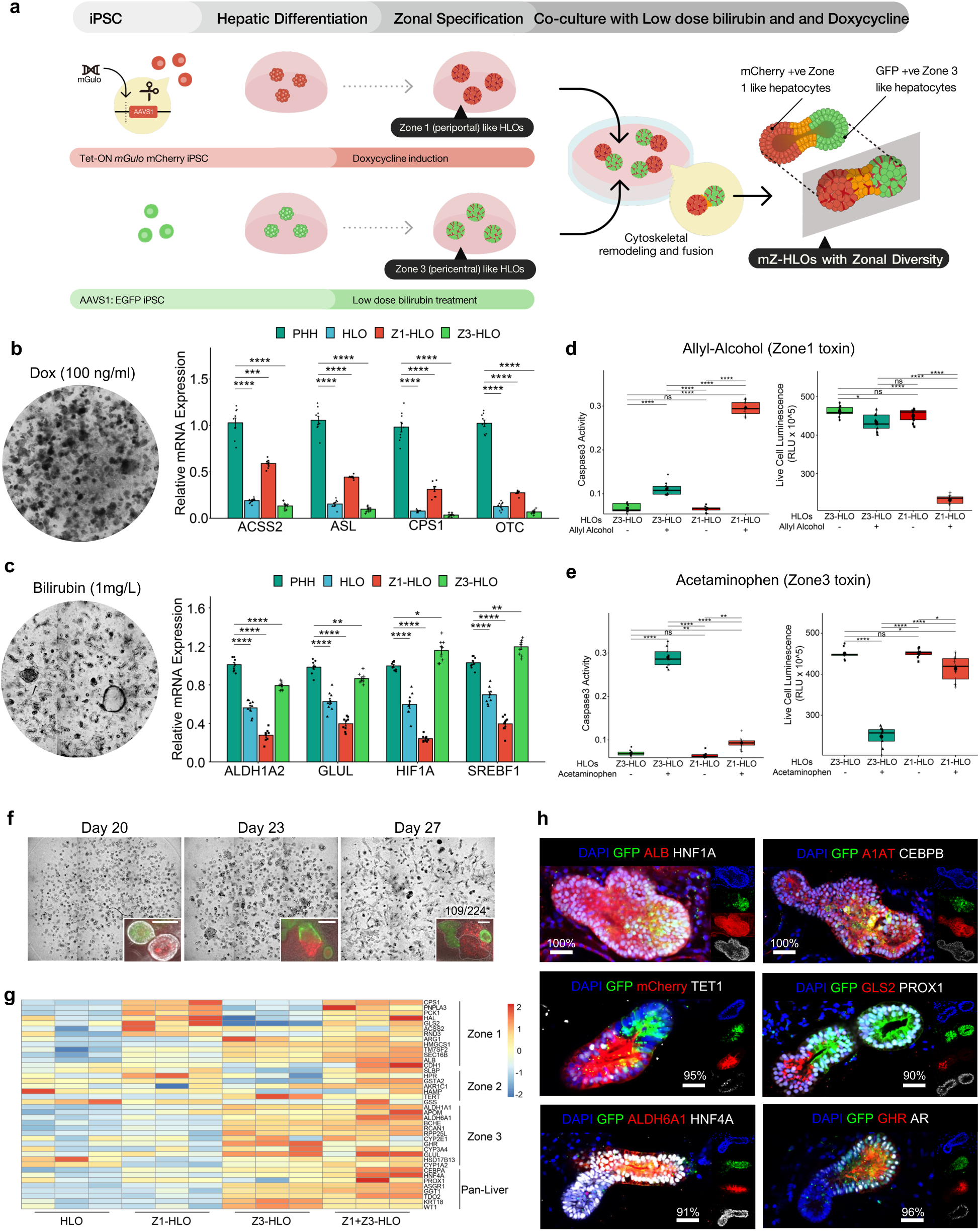
Engineering PSC-derived human liver organoids with multizonal hepatocyte features. **a**,Schematic for development of mZ-HLOs from doxycycline induced Z1-HLOs and low dose bilirubin treated Z3-HLOs. **b**,Brightfield images of Dox (100 ng/ml) treated Z1-HLOs (left). RT-qPCR of *ACSS2, ASL, CPS1* and *OTC* gene for Z1-HLOs compared to Primary Human Hepatocytes (PHH), Z3- and control HLOs (right). (Data is mean ± SD, n = 9 independent experiments). **c**,Brightfield image of Z3-HLOs treated with low dose bilirubin (1mg/L) (left). RT-qPCR of *ALDH1A2, GLUL, HIF1A* and *SREBF1* gene for Z3-HLOs compared to PHH, Z1- and control HLOs (right). (Data is mean ± SD, n = 9 independent experiments). **d**, Caspase 3 activity assay in Z3-HLOs compared to Z1-HLOs after treatment with Zone 1 toxin, Allyl alcohol, compared to control (left). Cell viability assay in Z3-HLOs compared to Z1-HLOs after treatment with Zone 1 toxin, Allyl alcohol, compared to control (right). (n = 9 independent experiments). **e**,Caspase 3 activity assay in Z3-HLOs compared to Z1-HLOs after treatment with Zone 3 toxin, Acetaminophen, compared to control (left). Cell viability assay in Z3-HLOs compared to Z1-HLOs after treatment with Zone 3 toxin, Acetaminophen, compared to control (right). (n = 9 independent experiments). **f**, Brightfield images (top) of progression of bilirubin induced fusion from Day 1 to Day 7. Fluorescent images of progression of organoid fusion from day 1 to day 7 when GFP+ Z3-HLOs were co-cultured with dox treated mCherry+ Z1-HLOs (inset). Scale bar indicates 200 µm. Numbers on the bar indicate the ratio of correctly fused organoids and total number of organoids. **g**, Heatmap of fused organoid compared to Z1-, Z3-, and control HLOs depicting expression of all zonal genes in the fused organoids. **h**, Immunofluorescence images of mZ-HLOs for pan liver markers: ALB, HNF4A, PROX1, HNF1A, A1AT, and CEBPB; Zone 1 markers: mCherry, TET1, and GLS2; Zone 3 markers: GHR, AR, and ALDH6A1. Scale bar indicates 200 µm. Numbers on the bar indicate the percentage of fused organoids that express dual and single positivity for the indicated antigen staining.

### Extracellular bilirubin specifies GLUL+ hepatocytes in Z3-HLOs

Parallelly, we treated a separate batch of HLOs expressing constitutive GFP with bilirubin at around Day 20 and found that 1 mg/L concentration enabled the greatest cellular survival in the HLOs (**Fig. 1a** and **Extended Data Fig. 2a and b**). Morphological analysis of the organoids revealed that they were more compact resulting in a smaller irregular lumen (**Fig. 1c, Extended Data Fig. 2c and d**). Aside from these morphological changes, the bilirubin treated human liver organoid expressed more zone 3 genes such as *ALDH6A1 (Aldehyde dehydrogenase 6A1), OATP2 (Organic anion transporter polypeptide 2),* and *GHR (Growth hormone receptor),* hereafter defined as Z3-HLOs (**Extended Data Fig. 2e**). Additionally, the Z3-HLOs expressed zone 3 specific *ALDH1A2 (Aldehyde dehydrogenase 1A2), GLUL, HIF1A,* and *SREBF1 (Sterol regulatory element-binding transcription factor 1)*, which were higher when compared to control HLOs and similar to PHH expression (**Fig. 1c**). Moreover, these Z3-HLOs exhibited higher CYP3A4 and CYP1A2 activity when compared to control, Z1-HLOs, and PHH (**Extended Data Fig. 2f**). Finally, the Z3-HLOs expressed pericentral specific GLUL and NR3C1 proteins (**Extended Data Fig. 2g**). Taken together, the data suggests that exposure to 1 mg/L bilirubin facilitates the differentiation into GLUL+ pericentral hepatocyte like population.

To further profile Z1- and Z3-HLOs, RNAseq were conducted over the respective HLOs. First, pan-hepatocyte marker genes such as *A1AT, HNF4A, and CEBPA* are consistently expressed in both Z1- and Z3-HLO. The Z3-HLOs, however, express more pericentral specific genes such as *GHR, BCHE (Butyrylcholinesterase)*, and *RCAN1 (Regulator of calcineurin 1)*, while Z1-HLOs express *ACSS2, SLBP (Stem-loop binding protein)*, and *RND3*, however they lack expression of widespread markers such as *ARG1* and interzonal markers such as *AKR1C1 (Aldo-keto reductase 1C1)* and *APOM (Apolipoprotein M)* (**Extended Data Fig. 2h**). Primary hepatocyte culture systems have been previously used to show that allyl alcohol and acetaminophen are toxic to zone 1 and zone 3 respectively due to differential alcohol and drug metabolism activity ^14^. By leveraging this pharmacological approach, we observed that the Z1-HLOs were sensitive to the zone 1 toxin, allyl alcohol, while Z3-HLOs were sensitive to the zone 3 toxin, acetaminophen as evidenced by the increased Caspase 3 activity (0.29 and 0.27 vs 0.02 relative activity) and reduced cell viability (225 and 250 vs 450 RLUX10^5^) (**Fig. 1d and e**). These indicated the acquisition of zone 1 or 3-like identity in HLO under specific inductive conditions.

### Multi-Zonal Human Liver Organoids (mZ-HLOs) via assembly of Z1- and Z3-HLOs

Next, we sought to generate a compartmentalized dual zonal organoid system (**Fig. 1a**). Upon further evaluation, we observed that the continuous bilirubin and Dox treatment induced the HLOs to self-assemble together, which is dependent on seeded organoid density under the persistent bilirubin exposure (**Extended Data Fig. 3a and b**). Real time imaging over time showed that individual organoids fuse together by the interaction of cytoskeletal proteins while maintaining a continuous lumen (**Extended Data Fig. 3c**). We observed that with continuous 1 mg/L bilirubin treatment the HLOs tend to fuse together indicated by the increase in mean segment length (**Extended Data Fig. 3d**). Approximately, 75% of this bilirubin treated organoids underwent fusion accompanied by increased Notch activity (Extended Data Fig. 3e and f). With inhibition of Notch signaling and Ezrin, the organoids failed to fuse together as observed over the course of 7 days suggesting a role of Notch and Ezrin in cytoskeletal rearrangements (**Extended Data Fig. 3f** and g). About 75% of the bilirubin treated organoids underwent fusion compared to <5% fusions when treated with DAPT or NSC668394. Thus, bilirubin induced Notch and Ezrin signaling activate cytoskeletal rearrangement to induce fusion. Fused organoids expand canalicular connectivity as was observed in a fluorescently labelled bile acid analogue transport assay (**Extended Data Fig. 3h**). Quantification of self-assembly efficiency indicated the preferential fusion efficiency in the cell line derived HLOs as follows: Z3-Z3 HLO (60%), Z3-Z1 HLO (35%), and Z1-Z1 HLO (5%) (**Extended Data Fig. 3i**).

We further characterized self-assembled dual organoid generated from the bilirubin treated Z3-HLOs and the dox treated Z1-HLOs (**Fig. 1f**). The fused organoids had more comprehensive expression of Zone 1, 2, 3 and pan-liver markers (*ACSS2, ALDH6A1, AKR1C1*, and *HNF4A*) by bulk RNAseq (**Fig. 1g**). Over the course of 10 days of continuous dual treatment, the most fused organoids generated structures carrying bilirubin in the lumen (**Extended Data Fig. 4a**). These putative multi-zonal organoids (mZ-HLOs) had a GFP+ zone 1 and a mCherry+ zone 3 side while expressing pan liver markers: A1AT (Alpha-1 antitrypsin) and PROX1 (Prospero homeobox 1) along the entire axis (**Extended Data Fig. 4b**). The mZ-HLOs also expressed pan hepatocyte markers: ALB, HNF1A, A1AT, CEBPB (CCAAT/enhancer-binding protein beta), PROX1, HNF4A, and TUBA1A (Tubulin alpha-1A) (**Fig. 1h**). They maintained a tubular structure indicated by the basal marker CTNNB1 and a continuous lumen indicated by the ZO-1 (Zonula occludens 1) apical marker (**Extended Data Fig. 4c**). Immunofluorescence also confirmed protein expression of three distinct regions within the newly generated mZ-HLOs: ARG1 (Arginase 1) positive region, TERT-positive region, and AHR (Aryl hydrocarbon receptor)- positive region, consistent with periportal, interzonal and pericentral zonal marker expression (**Extended Data Fig. 4c**). The other zone-specific liver markers such as apical MRP2 (Multidrug resistance-associated protein 2) and nuclear SLBP indicate the variable hepatic characters, while only a tiny proportion of the cells express the cholangiocyte marker CK7 (Keratin-7) (**Extended Data Fig. 4c**). Furthermore, the mCherry+ zone 1 side expressed zone 1 markers: TET1 and GLS2 (Glutaminase 2), while the GFP+ zone 3 side exhibited expression of zone 3 markers: ALDH6A1, GHR and AR (Androgen receptor) (**Fig. 1h**). These stains were consistent with the spatial patterning of TUBA1A, CK7, SLBP, GLS2, ALDH6A1, ARG1, TERT, AHR, and MRP2 expression in human neonatal liver tissue (**Extended Data Fig. 4d**).

### snRNAseq of mZ-HLOs reveal features of zonal hepatocytes

To detail the hepatic populations in the mZ-HLOs, we ran the samples through a single-nucleus (sn)RNAseq pipeline and retained 120,195 high-quality nuclei following stringent filtering for downstream analysis. The mZ-HLOs contained distinct populations that were like *SERPINA1*+ hepatocytes (55%), *KRT7*+ cholangiocytes (11%), *PECAM1*+ endothelial cells (12%), *LYZ*+ macrophages (7%), *COL1A1*+ stellate cells (7%), and *CD44*+ mesenchyme (8%) that exhibited discrete gene expression profiles (**Extended Data Fig. 5a and b**). We aimed to characterize the different hepatocyte identity, illuminate their functionally relevant gene sets, and explore their developmental trajectory using the snRNAseq dataset. Using unsupervised clustering and past hepatocyte expression profiles, we identified the Pericentral (28%) and periportal (24%) hepatocytes, as well as a hepatoblast population (30%) and two distinct interzonal hepatocyte population (18%) (**Fig. 2a**). Hepatoblasts are the most immature population in mZ-HLO and enriched for fetal markers such as *AFP,* and other growth mitogenic markers such as *IGF2 (Insulin-like growth factor 2),* and *MAP2K2 (mitogen-activated protein kinase 2),* which regulate the growth and differentiation of the cells. Interzonal hepatocytes are known for expressing glutathione and DNA repair enzymes and as such express *TERT* and *GSS*. Finally, the periportal hepatocytes expressed *GLS2, CPS1, OTC, ACSS2* and *ARG1*, while the pericentral hepatocyte population expressed *GLUL, CYP2E1, HIF1A, ALDH1A2, ALDH6A1*, and *AR* (**Fig. 2b and Extended Data Fig. 5c-e**). *TAT, HAMP,* and *CYP3A4* are localized in the periportal, interzonal and pericentral hepatocyte populations as cross-referenced by primary liver spatial transcriptomic dataset (**Extended Data Fig. 6a, b, c, and Fig. 2a**). Cholangiocyte marker *KRT7* was expressed at a minimal level in the hepatoblast population only (**Fig. 2b**), while extensive expression of *TTR (Transthyretin)* and *SERPINA1* was observed throughout the populations (**Fig. 2b**).

**Fig.2.**
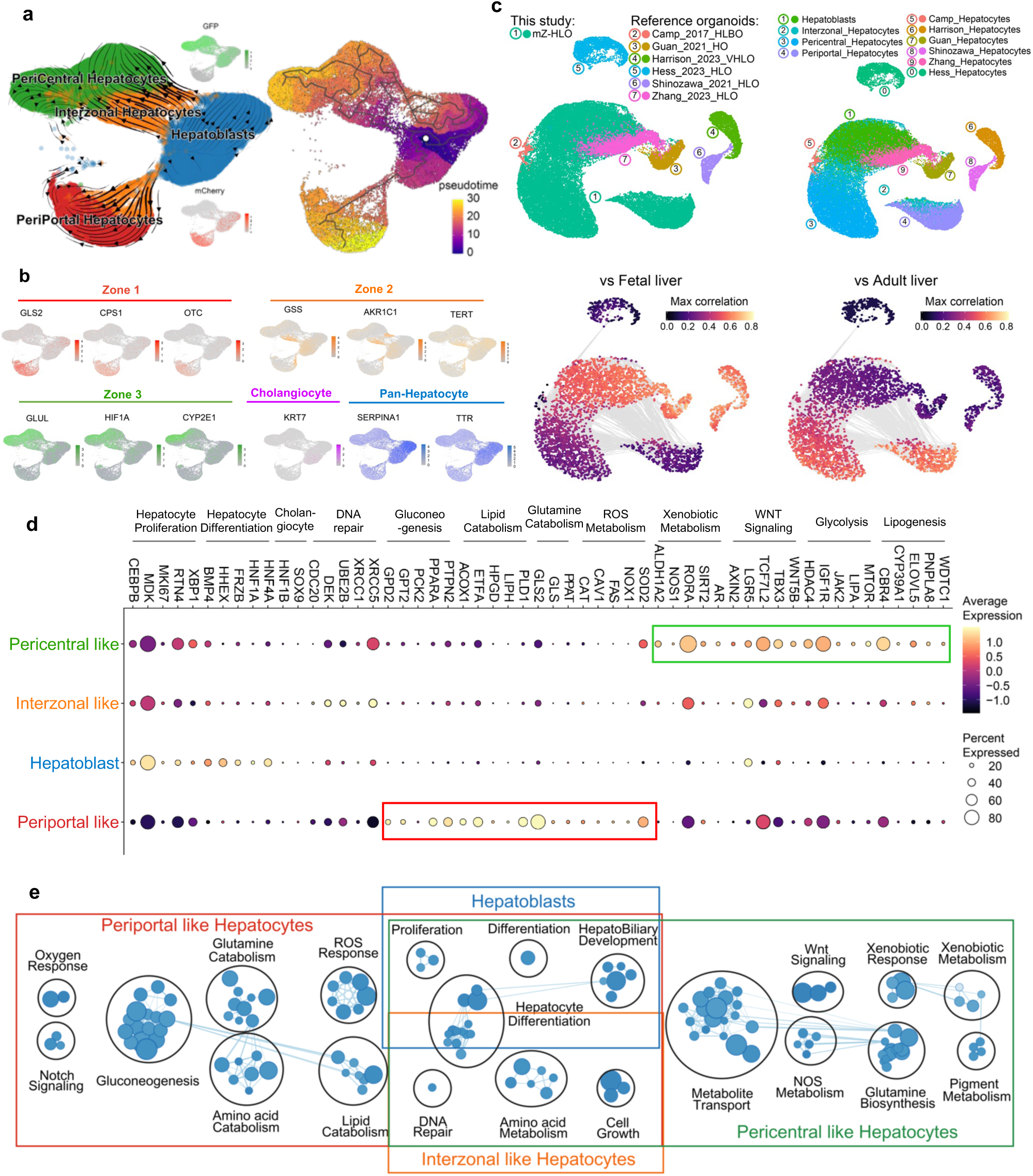
Single cell analysis of multi-zonal human liver organoids (mZ-HLO). **a**,UMAP plot with the major populations (Hepatoblasts, Interzonal like hepatocytes, Pericentral like hepatocytes, and Periportal like hepatocytes) of parenchymal nuclei in mZ-HLOs. Velocyto force field showing the average differentiation trajectories (velocity) for nuclei located in different parts of the UMAP plot (left). Pseudotime trajectory graph showing the differentiation trajectory for nuclei in the UMAP plot (right). The color represents the pseudotime development stage. **b**, Feature plots for pan liver makers: *TTR* and *SERPINA1*; Cholangiocyte marker: *KRT7*; Zone 1 marker: *mCherry, GLS2, CPS1* and *OTC*; Zone 2 marker: *GSS, TERT,* and *AKR1C1*; and Zone 3 maker: *GFP, GLUL, CYP2E1* and *HIF1A*. **c**, UMAPs for human hepatocytes from PSC-derived liver organoid cell atlas colored by organoid source and cell type. UMAPs displaying the maximum spearman correlation of fetal liver (left) and adult liver (right) dataset. **d**, Expression of genes related to zone specific functions in each population. The size of the circle indicates the percentage of nuclei in each population expressing each gene. The color represents the average expression level for the indicated gene. **e**, Pathway enrichment analysis examining which cellular pathways represented in the hepatoblast, pericentral, periportal, and interzonal hepatocyte populations. Circles (nodes) represent pathways, sized by the number of genes included in that pathway. Related pathways, indicated by light blue lines, are grouped into a theme (black circle) and labeled. Intra-pathway and inter-pathway relationships are shown in light blue and represent the number of genes shared between each pathway.

Additionally, we integrated our mZ-HLO dataset with multiple previously published adult and fetal hepatocyte snRNAseq datasets ^37–40^. Upon integration with the Andrews *et al.* dataset, we found a significant overlap of mZ-HLO-derived periportal, interzonal and pericentral hepatocyte populations with their primary counterparts, and the hepatoblast population clearly separated out into a different cluster (**Extended Data Fig. 6d**). Moreover, our periportal population expressed *GLS2, CDH1, CPS1* and *TET1* same as theirs, and our pericentral population also expressed *ALDH1A2, GHR, GLUL,* and *HIF1A* similar to theirs, while the interzonal profile was also alike (**Extended Data Fig. 6e**). Looking at global integration with all 4 datasets, there seems to be considerable overlaps particularly in terms of zonal hepatocytes (**Extended Data Fig. 6f**). There were modest overlaps between some of the adult liver zonal populations and the mZ-HLO zonal hepatocytes particularly in both pericentral and periportal hepatocytes in terms of *GLUL* and *GLS2* expression (**Extended Data Fig. 6g**). The mZ-HLOs also contain a hepatoblast population which is found in fetal liver along with other zonal features.

Furthermore, to benchmark our mZ-HLO model against existing models, we constructed an integrated dataset using a collection of 8 publicly available human PSC-derived liver organoid datasets using neighborhood graph correlation mapping (**Fig. 2c**) ^36,40–47^. With respect to the status of hepatocytes, the mZ- HLO showed a higher correlation with adult primary liver tissue (**Fig. 2c**), albeit the hepatoblast population exhibited higher correlation with fetal liver tissue.

We next queried the lineage progression into multizonal populations using RNA velocity and pseudotime analysis which is based on the kinetics of the splicing rate of mRNA and the expression of each gene (**Fig. 2a, Extended Data Fig. 7a and b**). Both methods predicted that the zonal hepatocytes originated from hepatoblasts through the interzonal hepatocyte population (**Fig. 2a and Extended Data Fig. 7c**). Finally, with the goal to predict the function of these individual populations we applied Gene Set Enrichment Analysis (GSEA) to identify the functional pathways and profiles of each set (**Fig. 2d and e**). Hepatoblast and interzonal hepatocyte populations were predicted to be mostly involved in hepatocyte proliferation and differentiation with interzonal cells more involved in DNA repair mechanisms. Periportal and pericentral hepatocytes were however vastly different. The periportal cells were involved in gluconeogenesis, lipid and glutamine catabolism, ROS and oxygen response, and was enriched for Notch signaling. On the other hand, Xenobiotic and pigment metabolism, Glutamine biosynthesis, and Wnt signaling were more enriched in the pericentral population (**Fig. 2d and e**). Thus, our self-assembled organoids harbor cells expressing zonal hepatocyte like markers.

### Differential EP300 regulation executes zonal transcriptional programs in mZ-HLOs

To understand the differential transcriptional mechanisms leading to zonation, we investigated the epigenetic landscape of the mZ-HLOs. EP300 (E1A-associated protein p300) is a histone acetyltransferase that acetylates enhancer regions and activates transcription leading to hepatoblast differentiation ^48,49^. EP300 marks poised and active enhancers and activates expression of zonal genes ^50–52^. EP300 ChIPseq analysis revealed that mZ-HLOs had increased binding of EP300 at putative enhancer sites (7875 binding sites) compared to control and singly treated HLOs (2852, 4891 and 5219 binding sites) (**Extended Data Fig. 8a**). Pan liver markers such as *HNF4A* and *CTNNB1 (β-catenin)* had EP300 peaks upstream of the TSS (Transcription Start Site) in all the samples similar to the PHH dataset in Smith *et al.* (**Extended Data Fig. 8b and c**) ^53^. However, many zone-specific genes such as *ACSS2* (zone 1), *ALDH6A1* (zone 3), and *HPR (Haptoglobin-related protein)* (zone 2) were recapitulated in mZ-HLOs while Z1- and Z3-HLOs had active enhancers upstream of only their respective zonal genes (Fig. 3a, b, and c). Furthermore, this pattern was again observed for *SLBP* (zone 1), *GHR* (zone 3), and *AKR1C1* (zone 2) as well (**Extended Data Fig. 8d, e, and f**). GSEA analysis of the peaks revealed enrichment for mixed zonal processes in the mZ-HLOs, while the Z1- and Z3-HLOs had zone specific biological process enrichment (**Fig. 3d, Extended Data Fig. 8g and h**). Motif analysis of the peaks also revealed that EP300 bound peaks were enriched for genes (*HNF1AI, HNF4A,* and *TBX2*) that regulate functional differentiation of hepatocytes (**Fig. 3e**). We then integrated our RNAseq dataset with the ChIPseq dataset to look at the regulation of upregulated RNA in each condition (**Fig. 3f**). The Z3-HLOs exhibited *HIF1A*, *TBX3 (T-box transcription factor TBX3)*, and *AHR* as the top motifs co-occupied by EP300, while Z1-HLOs revealed *TET1*, *NRF1 (Nuclear respiratory factor 1)*, and *TFEB (Transcription factor EB)* as the top motifs (**Fig.3f**). We further found that Bobcat339, a TET inhibitor, caused downregulation of *ACSS2* and *CPS1* in the Z1- and mZ-HLOs. While inhibition of HIF1A with KC7F2, a HIF inhibitor, downregulated *ALDH6A1* and *GLUL* (zone 3) expression in the mZ- and the Z3-HLOs (**Fig. 3g**). Together, these results suggested that EP300 co-ordinates with other TFs such as *HIF1A* and *TET1* to mediate the preferential activation and repression of zone-specific programs (**Fig. 3h**).

**Fig.3.**
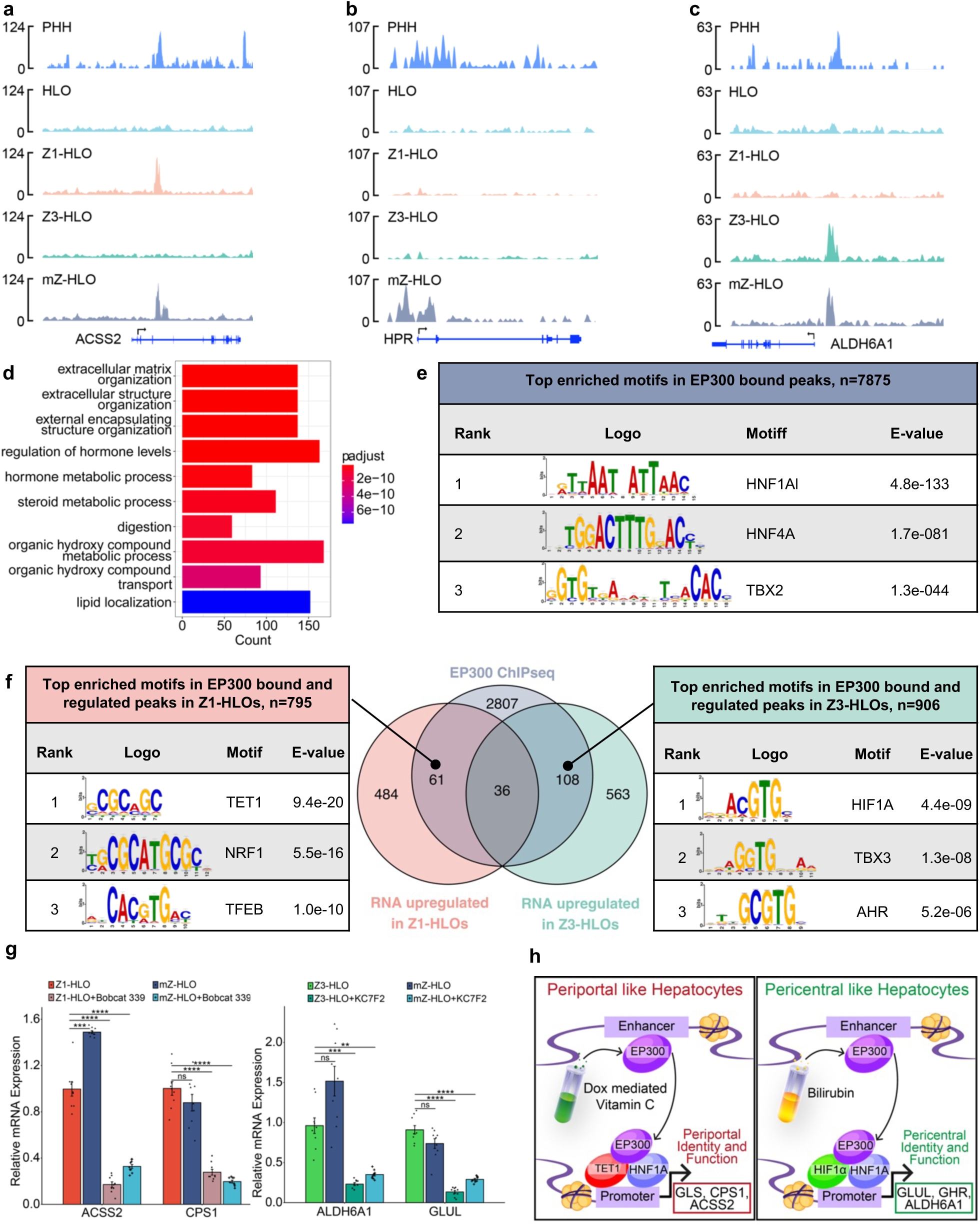
Bilirubin and Ascorbate regulate EP300 differentially in a spatial manner to evoke zonation. **a**, Genome browser view of ACSS2 (zone 1 gene) showing the EP300 ChIPseq peak. **b**,Genome browser view of HPR (zone 2 gene) showing the EP300 ChIPseq peak. **c**, Genome browser view of ALDH6A1 (zone 3 gene) showing the EP300 ChIPseq peak. **d**, Top 10 upregulated Gene Ontology terms (Biological Process) for the gene regulated bound by EP300 in the mZ-HLOs. **e**, Motif enrichment analysis of EP300 bound peaks analyzed by MEME-ChIP. **f**, Venn diagram depicting the intersection between EP300 bound peaks and upregulated genes obtained from RNAseq in Z1- and Z3-HLOs linked to motif enrichment analysis of EP300 bound peaks for upregulated genes in dox treated Z1-HLOs and in bilirubin treated Z3-HLOs analyzed by MEME-ChIP. **g**, RT-qPCR of *ACSS2,* and *CPS1* gene for Z1-HLOs compared to mZ-HLOs with and without treatment with Bobcat 339 (TET inhibitor) (left). RT-qPCR of *ALDH6A1,* and *GLUL* gene for Z3- HLOs compared to mZ-HLOs with and without treatment with KC7F2 (HIF1A inhibitor) (right). (Data is mean ± SD, n = 9 independent experiments). **h**, Schematic for bilirubin and ascorbate mediated distinct epigenetic regulation leading to differential gene expression.

For *in vivo* verification of the role of EP300 in zonal hepatocyte development, we injected Ad-shp300 into ∼P0-P1 rat pups via the retro-orbital route (**Extended Data Fig. 9a**). The *p300* shRNA mediated EP300 silencing resulted in impaired liver development where the hepatocytes underwent over proliferation and under differentiation causing portal and central vein ambiguity, *i.e.*, zonal impairment. This was further elucidated in the disrupted porto-central polarity mediated by EP300 silencing as evident in the aberrant expression of PROX1, ARG1, and GLUL. Here either PROX1 nuclear specificity or expression was completely lost in some hepatic cells, while for ARG1 and GLUL the zone-specific expression was altered to spotty non-zone-specific expression (**Extended Data Fig. 9a and b**). Furthermore, we compared our Z1- and Z3-HLO gene expression profile with size fractionated freshly isolated primary human hepatocytes ^54^. The Z1-HLOs had similar expression of *ACSS2, ALB, ASL, CPS1,* and *OTC* when compared to < 20 μm periportal hepatocytes (H20) (**Extended Data Fig. 9c**). On the other hand, the Z3-HLOs expressed *ALDH1A2, ALDH6A1, GLUL, HIF1A,* and *SREBF1* similar to the < 40 μm pericentral hepatocytes (H40) (**Extended Data Fig. 9d**). Moreover, when we looked at the ChIP-reChIP-PCR and ChIP-reChIP-qPCR, we found that H20 had increased binding of TET1 upstream of ACSS2 (1 vs 0.45-fold difference), while H40 had increased binding of HIF1A upstream of ALDH6A1 (1 vs 0.2-fold difference) similar to our Z1- and Z3-HLO respectively (**Extended Data Fig. 9e and f**).

### Zone specific programs work in concert for interzonal dependent nitrogen, lipid, and glucose handling in addition to proliferation in mZ-HLOs

Nitrogen metabolism in the liver occurs across multiple zones where ammonia is removed in the form of urea or glutamine ^5^. The urea cycle and glutathione metabolism pathways are intricately linked and required for proper nitrogen handling ^5^. Since we developed an organoid model with dual zonal functionality, we tested whether the mZ-HLOs could handle nitrogen metabolism that spans across multiple zones. The mZ-HLOs express urea cycle genes such as *CPS1, OTC,* and *ARG1*, while expressing detoxification genes such as *GSTA2 (Glutathione S-transferase A2), ALDH1A2,* and *GLUL* as well in response to 10mM NH_4_CL when compared to PHH and control HLOs (**Fig. 4a, b, and c**). This allows the mZ-HLOs to synthesize the highest levels of glutathione which can be inhibited by buthionine sulphoximine (BSO) (Fig. 4d). The mZ-HLOs therefore metabolize the highest amount of ammonia while BSO inhibits ammonia removal (**Fig. 4e**). This is reflected in the high urea levels detect in mZ-HLOs and the loss of urea production with BSO treatment (**Fig. 4f**). However, although urea cycle was inhibited with BSO treatment, ammonia levels were appreciably lower in the mZ-HLOs. We found that the mZ-HLOs had significant glutathione S-transferase activity and glutamine synthesis (**Fig. 4g and h**). This allowed the mZ-HLOs to upregulate glutamine synthesis to remove excess ammonia even in the presence of BSO treatment (**Fig. 4i**). Therefore, the mZ-HLOs are not only able to maintain dual zonal (zone1 and 3) functionality but also interzonal (zone2) dependent functions that are native to the nitrogen metabolism in the human liver as indicated by the BSO perturbation of interzonal functionality.

**Fig.4.**
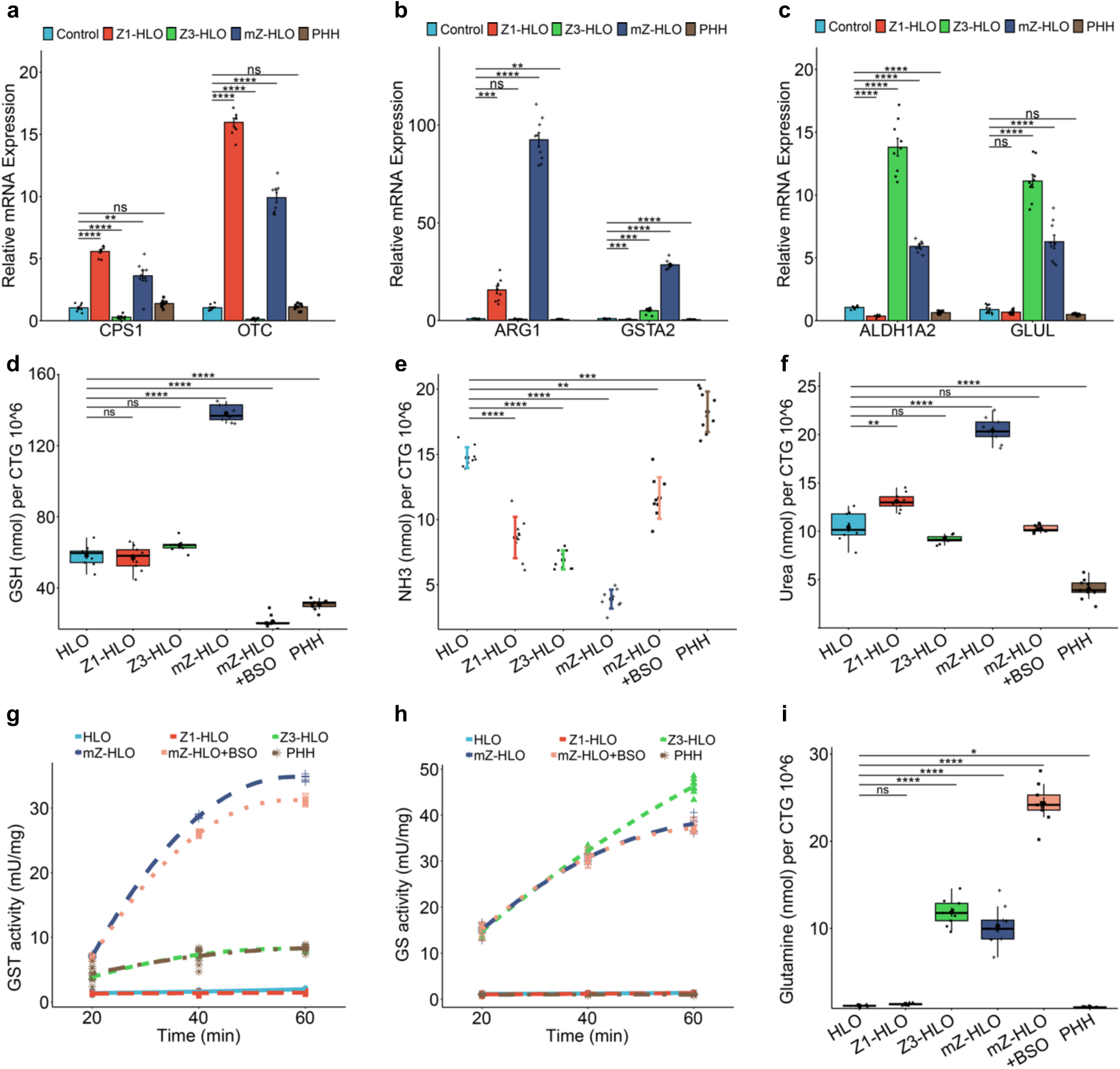
Interzonal dependent nitrogen handling in mZ-HLOs. **a**, RT-qPCR of *OTC* and *CPS1* (zone 1) gene for mZ-HLOs compared to Z1-, Z3-, control HLOs, and PHH in response to 10mM NH_4_CL (Data is mean ± SD, n = 9 independent experiments). **b**, RT-qPCR of *ARG1 (zone 1+2)* and *GSTA2* (zone 2) gene for mZ-HLOs compared to Z1-, Z3-, control HLOs, and PHH in response to 10mM NH_4_CL (Data is mean ± SD, n = 9 independent experiments). **c**, RT-qPCR of *ALDH1A2* and *GLUL* (zone 3) gene for mZ-HLOs compared to Z1-, Z3-, control HLOs, and PHH in response to 10mM NH_4_CL (Data is mean ± SD, n = 9 independent experiments). **d**, Glutathione assay for mZ-HLOs with and without BSO treatment compared to Z1-, Z3-, control HLOs, and PHH (n = 9 independent experiments). **e**, Ammonia assay for mZ-HLOs with and without BSO treatment compared to Z1-, Z3-, control HLOs, and PHH (n = 9 independent experiments). **f**, Urea assay for mZ-HLOs with and without BSO treatment compared to Z1-, Z3-, control HLOs, and PHH (n = 9 independent experiments). **g**, Glutathione S-Transferase assay for mZ-HLOs with and without BSO treatment compared to Z1-, Z3-, control HLOs, and PHH (n = 9 independent experiments). **h**, Glutamine synthetase activity assay for mZ-HLOs with and without BSO treatment compared to Z1-, Z3-, control HLOs, and PHH (n = 9 independent experiments). **i**, Glutamine assay for mZ-HLOs with and without BSO treatment compared to Z1-, Z3-, control HLOs, and PHH (n = 9 independent experiments).

Furthermore, since lipid and glucose metabolism, which are intrinsically linked, are maintained differentially by the zonal hepatocytes, we wanted to see whether the mZ-HLOs exhibit this metabolic balance ^2,55^. Consequently, we performed a triglyceride assay which showed that mZ-HLOs had intermediate levels of triglyceride (4.5 nmol) when compared to Z1- (2.5 nmol) and Z3-HLOs (5.5 nmol). Firsocostat (Acetyl-CoA Carboxylase inhibitor) treatment resulted in inhibition of lipid biosynthesis and the lowest levels of triglyceride due to the high lipase activity of the mZ-HLOs (**Extended Data Fig. 10a and b**). On the other hand, for glucose metabolism we looked at the glucose levels where mZ-HLOs had intermediate levels of glucose (23 nmol) as well compared to other zonal HLOs. FBPi (Fructose 1,6-bisphosphatase inhibitor) treatment inhibited the gluconeogenetic process resulting in lower levels of glucose due to enhanced glycolytic activity of the mZ- HLOs (**Extended Data Fig. 10c and d**).

Finally, we sought to test the zone-specific regenerative potential of the mZ-HLOs upon transient treatment with Allyl Alcohol (Zone 1 toxin) and Acetaminophen (Zone 3 toxin) (**Extended Data Fig. 11a**). We found that with the allyl alcohol, the GFP –ve regions had positive expression of Ki-67 starting at the intermediate region. Conversely, the acetaminophen transient treatment resulted in Ki-67 expression primarily in the GFP +ve regions (**Extended Data Fig. 11b and c**). This resulted in elongation of the GFP -ve ARG1 region and the GFP +ve GLUL region following treatment of allyl alcohol and acetaminophen respectively (**Extended Data Fig. 11b, d**). Furthermore, the Z1-HLO exhibited zone 3 features as evident in switching of CPS1 and TET1 expression to GLUL and NR3C1 expression at Day 25 following doxycycline withdrawal and persistent bilirubin treatment starting at Day 20 when the Z1 features started to appear (**Extended Data Fig. 11e**). This switch was also observed in reduced expression of ACSS2 and increased expression of ALDH1A2 following doxycycline withdrawal and bilirubin treatment (**Extended Data Fig. 11f**).

### Orthotopic transplantation of mZ-HLOs alleviates symptoms in bile duct ligated rats

To determine multi-zonal functionality *in vivo*, we evaluated post-transplant mZ-HLO metabolic performance in ammonium and bilirubin removal relative to singular zonal HLO system. In choosing a model, we adopted Bile duct ligation (BDL) because BDL in rats exhibits hyperammonemia and hyperbilirubinemia, leading to the progression of total hepatic dysfunction ^56^. The ligation results in large accumulation of bilirubin and ammonia in the serum that accelerates liver injury ^57^. Since the mZ-HLOs exhibit dual zonal functionality, we explored whether mZ-HLOs can be used in an *in vivo* setting for the amelioration of disease. We transplanted the mZ- HLOs orthotopically on the liver in close proximity of the portal vein using fibrin glue as a scaffold in bile duct ligated *Il2rg*-deficient, *Rag1*-deficient rats and observed them for 30 days (**Fig. 5a**). The mZ-HLO transplanted rats had a higher survival rate (55.6%, p = 0.049) compared to Z3-HLO (33.3%), Z1-HLO (22.2%), and sham transplants (22.2%) (**Fig. 5b**). The transplanted rats exhibited approximately 350 ng/ml peak human albumin in their serum at Day 20 indicating functional engraftment (**Fig. 5c**). Moreover, the transplanted mZ-HLOs were observed to invade into the hepatic parenchyma and retained their structure as indicated by the TUBA1A and ASGR1 stain (**Extended Data Fig. 12a and b**). These partially integrated mZ-HLOs retained their mCherry and GFP expression which also expressed ARG1 and GLUL (**Extended Data Fig. 12c**). Comparatively the Z1-HLO integrated into the parenchyma of the liver more efficiently when transplanted through the portal vein route, whereas the Z3-HLOs integrated better near the central vein when transplanted via the inferior vena cava route (**Extended Data Fig. 12d**). Subsequently, we observed reduction in levels of the elevated bilirubin (down from approximately 13 to 6.5 mg/L) and ammonia (down from approximately 160 to 70 μM) at Day 20 in the rat serum (**Fig. 5d and e**). Overall, our data suggests that the transplantation of the multizonal mZ- HLOs offers better metabolic performance in removing excess ammonia and bilirubin than non-multi-zonal tissue transplant.

**Fig.5.**
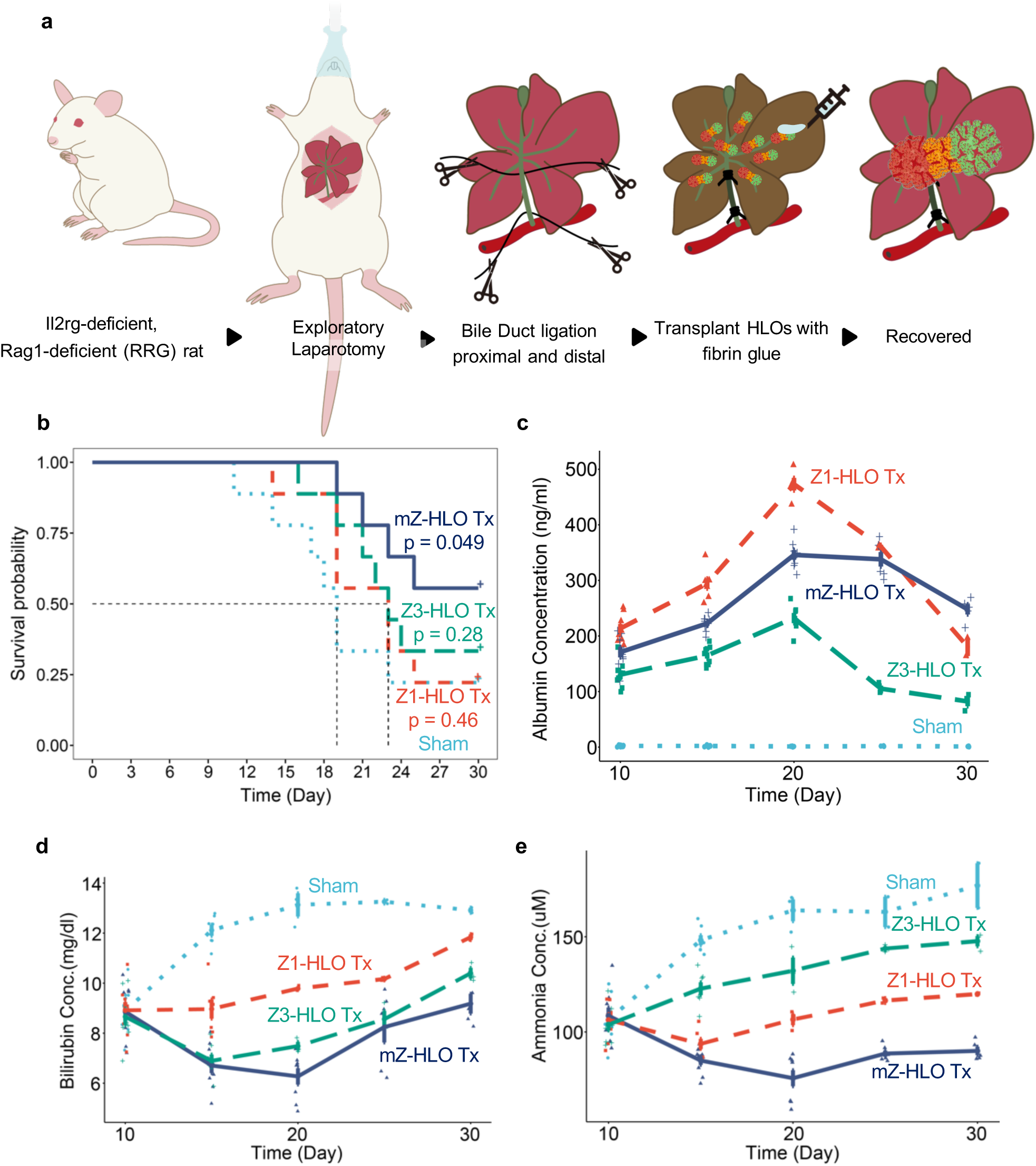
Orthotopic transplantation of mZ-HLOs improves multiple hepatocyte functions. **a**, Schematic for orthotopic transplantation of HLOs in bile duct ligated immunocompromised RRG rats. **b**, Kaplan-Meier survival curve for Z1-, Z3-, and mZ-HLO transplanted bile duct ligated RRG rats compared to sham. **c**, Human Albumin ELISA on blood serum collected from rats at different time points after transplantation. (n = 9 independent experiments). **d**, Bilirubin assay on blood serum collected from rats at different time points after transplantation. (n = 9 independent experiments). **e**, Ammonia assay on blood serum collected from rats at different time points after transplantation. (n = 9 independent experiments).

## Discussion

Handling both the synthesis and degradation of a multitude of metabolites requires precise compartmentalization, a feature that is achieved in the liver by a process called zonation ^58^. In zonation, the hepatocytes are specialized into periportal hepatocytes (Zone 1) located near the portal vein, pericentral hepatocytes (Zone 3) near the central vein, and a small population of interzonal hepatocytes in the intermediate region ^11^. The periportal hepatocyte develop in an oxygen and nutrient rich environment^13^, and vice versa for pericentral hepatocytes. Despite several attempts using primary and stem cell-derived hepatocytes, the emergence of multi-zone dependent functions in existing human hepatic tissue models has been unclear ^59,60^. Here we have developed a new platform for studying the development and zonal functionality using human liver organoids from pluripotent stem cells. Our data suggests that bilirubin and ascorbate are able to drive epigenetic and transcriptional programs towards the development of zonal hepatocyte identity. The mZ-HLOs are equipped with the urea cycle, glutathione, and glutamine metabolic functions to exert concerted efforts to control nitrogen availability as seen *in vivo*. Additionally, our mZ-HLOs are capable of handling the intricate symphony associated with metabolism of lipids and glucose as maintained by the native zonal liver. As the system we have developed is tractable and manipulable, mZ-HLO system paves a way for the study of development and disease affectingthe spatially and functionally divergent hepatocyte subpopulations found in humans.

Emerging single cell genomics approach revealed the high-resolution signatures that define zonation pattern in the liver ^39,61^. With hindsight, we opted for snRNAseq to characterize iPSC derived epithelial components in organoids ^62^. Our snRNAseq dataset showed divergent parenchymal populations including hepatoblasts, interzonal, pericentral and periportal hepatocyte-like cells, which were annotated based on the aforementioned datasets and well-known genetic markers. We also performed a comprehensive pathway analysis to determine the functional aspects of the individual populations. Moreover, we find that when compared to primary liver snRNAseq dataset, the zonal hepatocyte population are highly concordant with subpopulations found in mZ-HLO, with the exception of the hepatoblast population. The genes and pathways activated in zonal subpopulations are largely in agreement with existing knowledge related to specialized molecular markers: however, few discrepancies are noted such as lower expression of *TTR, CEBPB, APOA1* and *ARG1* when compared to publicly available primary liver-derived datasets. The discrepancy is possibly due to immaturity or differences between snRNAseq and scRNAseq. Interestingly, two different unsupervised trajectory analysis generate insights around the hepatoblast lineage progression to the zonal hepatocytes. Our developmental lineage predictions supported the assertion made by He et al. that zonal hepatocytes originate from hepatoblasts after differentiation through the interzonal hepatocyte fates ^63^. However, this trajectory model of our mZ-HLOs only depicts the development trajectory of only early zonal liver development. Due to the artificial nature of our mZ-HLOs, our model does not show signs of transdifferentiation beyond a certain time period, whereas adult hepatocytes have been reported to transdifferentiate into different zonal hepatocytes ^64^. Regardless, new evidence suggests that interzonal hepatocytes are the primary source of other zonal hepatocytes during liver regeneration which is very similar to our mZ-HLO model ^61,63^. Therefore, significant advances need to be made to address this trans- differentiation and regenerative capability. Collectively multi-zonal organoids together with single cell genomics will provide a framework to study the developmental divergence of hepatocytes in humans.

The epigenetic landscape determines the differentiation into zonal hepatocytes in the liver ^65^. EP300, a histone acetyltransferase, is one such epigenetic modifier that acetylates enhancer regions and activates transcription leading to hepatoblast differentiation ^48,49^. EP300 marks poised and active enhancers and activates expression of zonal genes ^50–52^. For instance, periportal metabolic functions such as gluconeogenesis and beta-oxidation of fatty acids are known to be activated by EP300 ^66,67^. On the other hand, EP300 is also known to contribute to the upregulation of genes involved in glycolysis and lipogenesis, functions specific to pericentral hepatocytes ^68,69^. Similarly, our data suggests that EP300 binds to enhancers upstream of zonal genes such as *ALDH6A1, ACSS2* and *HPR* in a context-dependent manner to differentially activate gene expression similar to previously published PHH studies. The integrated RNAseq and ChIPseq showed that the top targets were *HIF1A* and *TET1* in the dox treated Z1-HLOs and bilirubin treated Z3-HLOs respectively. Developmentally, the *Tet1* deletion impairs periportal identity and function in the liver, whilst repressing the pericentral characters regulated through Hedgehog signaling ^12,20^. Inversely, recent studies have demonstrated that bilirubin possesses signaling properties that can activate HIF and WNT signaling cascades ^30,70^. which are important for pericentral hepatocytes ^71^. Following such developmental patterns, mZ-HLOs exposed to inhibitors specific to HIF1A (KC7F2) and TET1 (Bobcat 339) lead to the loss of zonal identities in concordant with these publications ^12,20^. Moreover, a mouse siRNA model revealed that EP300 is integral to zonal liver development which further validated out results. Finally, freshly isolated size fractionated PHH showed that H20 (periportal hepatocytes) had a TET1 binding site upstream of ACSS2, while H40 (pericentral hepatocytes) had a HIF1A binding site upstream of ALDH6A1 identical to our mZ-HLOs. These results corroborate the developmental role of HIF1A and TET1 in executing zonal program, therefore future endeavors are also necessary to look into the other transcription factors including HNF1A, HNF4A and TBX2 using ChIPseq to evaluate their roles in development of the liver.

Hepatocyte transplantation has been used to treat liver diseases but the difficulty in obtaining compatible primary human hepatocytes makes this an impractical approach ^72^. Recently stem cell derived tissues have been used to treat a multitude of hepatic diseases ^73^. However, the most therapeutic proof-of-concept has aimed at correcting monogenic conditions by targeting one particular disease parameters. Our mZ-HLOs model is endowed with the zonal functionality that ameliorated multiple aspects of the liver failure *in vivo* in a bile duct ligated immunodeficient rat model. There was significant reduction in serum bilirubin and ammonia levels as detected in the rat, while serum protein albumin is secreted. All these data translate into an increased survival rate of the rats that received the mZ-HLO transplantation when compared to the sham. Interestingly, zone-1 or zone-3 primed organoid transplant only offer either hyperammonemia or hyperbilirubinemia improvements resulting in reduced survival benefit, indicating that the balanced delivery of multizonal hepatocytes is crucial for improving total liver dysfunctions. Furthermore, the mZ-HLOs integrated into the parenchyma of the rat liver while retaining their zonal characteristics as indicated by the expression of mCherry, GFP, ARG1, and GLUL. Finally, the Z1-HLOs showed a preferential integration near the portal vein, while the Z3-HLOs exhibited a slightly higher affinity for the central vein region. Overall, our data suggests that mZ-HLOs are able to engraft into the resident liver and maintain zonal-specific functionality and augment the survival rate in rodents following biliary duct ligation. Finally, mZ-HLOs may therefore be further explored as a tool which can recapitulate zonal identity and function critical in the study and treatment of hepatic disease.

## Methods

### Animals

All animal experiments were conducted with the approval of the Institutional Review Board (IRB) and Institutional Animal Care and Use Committee (IACUC) of the Cincinnati Children’s Hospital Medical Center. Adult *Il2rg*-deficient, *Rag1*-deficient RRG (SD/Crl) rats (breeding pairs, 9–12 weeks old) were housed in standard rat cages with paper bedding and maintained at a temperature of 20–24 °C and relative humidity of 45–55%, under a 12 h:12 h light:dark cycle. All animals had ad libitum access to dox chow before study. All animals were treated in accordance with the guidelines and regulations of the institution.

### mGulo editing

The murine Gulo (L-gulonolactone oxidase) cDNA sequence was retrieved from NCBI. The 5′ linker and Kozak sequence were added to the start of the sequence, with HA tags to the end of the sequence. Additionally, a P2A-mCherry was added after the HA tag and a 3′ linker to the very end. The custom gene was then synthesized and cloned into the pAAVS1-NDi-CRISPRi (Gen1) PCSF#117 vector using the restriction sites AflII and AgeI. The vector has a TetON system and a Neo^r^ selectable marker was then inserted using the Gateway technology.

### mGulo iPSC generation and general iPSCs maintenance

Experiments using iPSCs were approved by the Ethics Committees of Cincinnati Children’s Hospital Medical Center. The 1383D6 and RCL-BC iPSC used in this study was kindly provided by CiRA, Kyoto University. The iPSCs 72.3, and 72.3-GFP were obtained from patient foreskin fibroblasts and reprogrammed into iPSC by Cincinnati Children’s Hospital Medical Center pluripotent stem cell core. The PCSF#117 vector with the modified *mGulo* sequence was then inserted into the AAVS1 locus of the 72.3 iPSC cell line using a lentiviral mediated CRISPR/Cas9. The correct clones were then selected using G418. The surviving clones were then verified for correct insertion, random insertion and copy number using PCR, and verified by DNA sequencing. The iPSCs were then maintained on Laminin iMatrix-511 Silk (REPROCELL USA Inc.) coated cell culture plates and maintained with StemFit Basic04 Complete Type (Ajinomoto Company) media with Y-27632 (Stem Cell Technologies). The cells were passaged every 7 days with Accutase (Sigma-Aldrich) until passage 40 (p40).

### Organoid generation

The p40 cells were plated on a 24 well plate coated with Laminin iMatrix-511 Silk at a density of 2×10^5^ cells/well and maintained with Stemfit media with Y-27632. On Day 2, the media was replaced with fresh Stemfit. The following day, the cells were treated with RPMI 1640 (Gibco) media mixed with Activin A (Shenandoah Biotechnology) and BMP4 (R&D Systems) to generate definitive endoderm. On the 4th day, the media was replaced with RPMI, Activin A and 0.2% dFBS (HyClone) which was changed to 2% dFBS on day 5. From Day 6-8, the cells were fed with FGF-4 (Shenandoah Biotechnology) and CHIR99021 (PeproTech) in Adv. DMEM [Advanced DMEM/F-12 (Gibco) with B27 (Gibco), N2 (Gibco), 10mM HEPES (Gibco), 2mM L-glutamine (Gibco), and GA-1000 (Lonza)] to induce posterior foregut. On Day 9, the cells were dissociated into a single cell suspension using Accutase treatment. This single cell suspension was then mixed with 50% Matrigel and 50% EP media and plated as 50 ul drops in a 6-well plate. These cells were fed with EP media every 48 hrs for 4 days to generate organoids. These organoids were then treated with Adv. DMEM and 2 μM RA (Sigma-Aldrich) every 48 hrs for 4 days to specify the hepatic lineage. The organoids were then fed with HCM (Lonza), HGF (PeproTech), Oncostatin M (PeproTech) and Dexamethasone (Sigma-Aldrich) every 3-4 days to generate HLOs and passaged as necessary.

### Generating multi-Zonal Human Liver Organoids (mZ-HLOs) with zonal diversity

For generation of multi-zonated structures, the immature HLOs were treated with low conc. bilirubin (1 mg/L) in HCM on Day 20. The bilirubin treatment was maintained with every media change onwards by keeping the cells at 37°C in 5% CO_2_ with 95% air. The Z1-HLOs (Zone 1) were maintained with Dox starting at Day 17 and co-cultured on Day 22 with the bilirubin treated 72.3-GFP (GFP+) Z3-HLOs (Zone 3) in a 1:1 ratio at higher density *i.e.,* 2× the number of organoids with continuous bilirubin and Dox treatment in HCM to obtain chimeric organoids that had dual zonal characteristics. These mZ-HLOs and HLOs were visualized by using fluorescent microscopy BZ-X810 (Keyence, Osaka, Japan) and harvested for downstream analysis. The images were analyzed with Fiji (v1.53f) for morphometric and quantitative measurements.

### Live cell imaging and functional assay

For live imaging of organoids, the CellDiscoverer 7 (Zeiss) was used to image every 30 min for 7 days. Observing organoid fusion necessitated looking at the cytoskeleton, the HLOs were incubated with 2 drops/ml NucBlue (Hoechst 33342) (Invitrogen, R37605) and 1 μM Cytoskeleton Kit (SiR-Actin and SiR-Tubulin) (Cytoskeleton Inc., CYSC006) and imaged over 5 days. For functional assay of lipid transport, the HLOs were incubated with fresh media containing 50 nM CLF (Corning, 451041) before imaging them every 30 min for 2 days.

### RNA extraction, RT-qPCR, and RNA sequencing

RNA was isolated using the RNeasy mini kit (Qiagen, Hilden, Germany). Reverse transcription was carried out using the High-Capacity cDNA Reverse Transcription Kit for RT-PCR (Applied Biosystems) according to manufacturer’s protocol. qPCR was carried out using TaqMan gene expression master mix (Applied Biosystems) on a QuantStudio 5 Real-Time PCR System (Applied Biosystems). All the samples were amplified with TaqMan Gene Expression Assays and normalized with 18S rRNA Endogenous Control. Human primary hepatocytes (Lonza Catalog #: HUCPG and Sigma-Aldrich’s Catalog #: MTOXH1000) were used as PHH control. For RNA sequencing the service was outsourced to Novogene (USA), the extracted RNA quality was evaluated with an Agilent 2100 Bioanalyzer (Agilent). A sequence library was prepared using a TruSeq Stranded mRNA kit (Illumina) and sequenced using NovaSeq 6000 (Illumina). Reads were aligned to human genome assembly hg38 and quantified using the quasi-mapper Salmon (v1.8.0). Gene-expression analysis was performed using the R Bioconductor package DESeq2 (v1.36.0). The read count matrix was normalized by size factors, and a variance stabilizing transformation (VST) was applied to the normalized expression data. The data was visualized using clusterProfiler (v4.4.2) and pheatmap (v1.0.12) packages.

### ChiP-reChIP (-PCR, and -qPCR), ChIP-seq and analysis

ChIP experiments were performed using the High Sensitivity ChiP Kit (Abcam, ab185913). Briefly, organoids were fixed with PFA and whole chromatin was prepared and then sonicated to an optimal size of 300bp which was confirmed by gel electrophoresis. Chromatin was used for immunoprecipitation (IP) with either EP300 antibody (ab14984) or IgG1 isotype control. For the ChIP-reChIP, the ACSS2 (Abcam, ab133543) and ALDH6A1 (Abcam, ab12618) antibody was crosslinked to Protein A Dynabeads (Invitrogen, 10002D). The ChIP assay was then carried out on extracts from organoids as described above. At the end of the first ChIP, DNA was eluted with elution buffer supplemented with 10 mM DTT. The eluate was then diluted in 2 volumes of wash buffer supplemented with 1x Protease Inhibitor Cocktail and 1 mM DTT. The 2nd ChIP assay was then carried out as described above. For ChIP sequencing, the service was outsourced to MedGenome (USA), the quality of the DNA was analyzed with Qubit (Invitrogen) and TapeStation (Agilent). A sequence library was prepared using a NEB Next Ultra II DNA kit and sequenced using NovaSeq 6000 (Illumina). Reads were trimmed and quality-checked using TrimGalore (v0.6.6) and then aligned to hg38 using bwa (v0.7.17). The aligned files were filtered, sorted and indexed using SAMtools (v1.15.1), and unmapped and low quality (MAPQ<30) reads were discarded. The duplicates were then marked and removed with Picard (v2.27.3). For visualization, deepTools (v3.5.1) was used to generate BigWig files which were visualized using IGV (v2.13.0). Peaks were identified using MACS2 (v2.2.7.1) and annotated with ChIPseeker (v1.32.0) to generate BED and BEDgraph files for visualization with IGV. For differential binding analysis, DiffBind (v3.6.1) was used to call statistically significant differential peaks after normalization and differential regions were selected based on DESeq2 method FDR-corrected q-value of 0.05. Heatmap and profile plots were generated with EnrichedHeatmap (v1.26.0). The functional analyses of GO term and KEGG pathway were performed using clusterProfiler. De novo motif analysis was then carried out on centered 100 bp regions from the peak summits using MEME Suite (v5.4.1).

### snRNA-seq and analysis

For snRNA-seq, 25-30 mg samples were pulverized with liquid nitrogen and nuclei were prepared using Nuclei EZ Lysis buffer (NUC-101; Sigma-Aldrich) as described by Wu et al. with addition of 0.04% BSA/PBS in the final buffer ^62^. The nuclei were filtered through a 10 μm filter, sorted, and counted before the library was generated using the Chromium 3′ v3 GEM Kit (10x Genomics, CG000183RevC). Sequencing was performed by the CCHMC DNA Sequencing core using the NovaSeq 6000 (Illumina) sequencing platform with an S4 flow cell to obtain approximately 320 million reads per sample. The demultiplexing, barcode processing, gene counting, and aggregation were done and the fastq files were aligned to the GRCh38 human reference transcriptome using cellranger v7.0.1, alevin-fry v0.8.0, and starsolo v2.7.9a. to extract the UMI and nuclei barcodes. SoupX v1.6.0 was used to remove ambient RNA and other technical artifacts from the count matrices. The UMIs were quantified per-gene and per-nuclei for normalization. The dataset was then analyzed using Seurat v4.2.0 in RStudio v4.1.1. Quality control was then carried out by using filtering parameters where nuclei with features less than 200 and greater than 4000 or more than 0.5 percentage mitochondrial genes were discarded. In the end, 45,223 parenchymal nuclei were isolated out from a total of 120,195 nuclei. The dataset was then normalized, and top 2000 highly variable genes were selected using the ‘VST’ method. The dataset was then scaled, and principal component analysis (PCA) was run for dimensional reduction. Elbow plots and JackStraw plots were then used to determine the number of PCs to be used. The nuclei were then clustered using Louvain algorithm and KNN. Uniform Manifold Approximation and Projection (UMAP) projections were then used to visualize the clusters. Nulcei clusters were annotated based on gene expression levels of known markers and markers detected in previously published datasets ^39,61^. GSEA analysis was then carried out using clusterProfiler v4.4.2; the rank files were then extracted to group highly related pathways to the specific clusters in Cytoscape v3.9.1. For integration, previously published dataset was merged with our dataset, normalized, integration features and anchors were computed, scaled, dimensions reduced, clusters created and finally projected as UMAP reductions. For trajectory analysis, monocle3 v1.2.9 was employed to convert the dataset to ‘cds’ objects, partitions were created, the trajectory graph was learnt in an unsupervised fashion, and finally the nulcei were ordered in psuedotime. Concurrently, the dataset was imported into scanpy v1.9.1 as ‘anndata’ and trajectory was analyzed using scvelo v0.2.4 by projecting the computed RNA velocity onto the previously generated UMAP reduction.

### Constructing and comparing an integrated dataset of published human liver organoids and a primary liver reference

For integration 8 different protocol-based human PSC-derived liver organoid and 4 primary adult and fetal datasets were collected according to the descriptions in the original publications ^36,41–45^. Briefly, we obtained available data (either raw FASTQ files, count matrices, H5AD, or Cell Ranger outputs such as “filtered_feature_bc_matrix” files) for each organoid from databases such as GEO, ArrayExpress, and the Human Cell Atlas (HCA). For FASTQ files, we used Cell Ranger to align and quantify the sequencing reads with the same parameters described in the original publication, generating UMI count data. Subsequent data processing was performed in Seurat using default settings. We curated metadata for all organoid data, including cell barcodes, sample names, cell type annotations, and cell cycle phase. We normalized and combined the public organoid data with our mZ-HLO data. Concurrently, we identified the top 3,000 variable genes from the primary liver data and applied these to the organoid dataset. Cell type annotations were based on the original publication and assigned into hepatocytes, hepatoblasts, endothelial cells, cholangiocytes, macrophage, mesenchyme, and stellate cells, which were added as new metadata. Using Seurat RPCA integration, we integrated the organoid data comprising 29,526 cells and the primary liver data comprising 8,656 cells. The same configurations were used to integrate the mZ-HLO dataset. After integration, we performed Louvain clustering and re-annotated cell types based on the expression of known marker genes. For further comparative analyses, we used the integrated organoid data as the query and the primary liver data as the reference. To benchmark our mZ-HLO model against existing models, we used the miloR and scrabbitr R packages ^46,47^ to compute neighborhood graphs, compare neighborhoods based on similar features, and map neighborhood comparison defined by k-NN graph using UMAP embeddings for primary adult and fetal liver dataset. The neighborhood correlations were computed using 3000 highly variable genes that were found in the highly variable genes in either adult or fetal primary liver compared as reference. The transcriptional similarity graph was computed using 30 dimensional nearest neighbors and UMAP embeddings of cells, while other parameters were implemented as default.

### Adenovirus mediated gene silencing of p300 in vivo

The BLOCK-iT adenoviral RNA interference expression system (Invitrogen, Carlsbad, California) was used to construct adenoviral short hairpin RNA (shRNA) for p300 and scrambled shRNA as previously described ^74^. Rat pups aged ∼ P0-P1 were then placed on a sterile heating pad, sanitized using isopropyl alcohol and iodine tincture to clean the skin surface. Finally, a 32G 1 inch needle was used to inject the adenoviruses (10 µg = 1.25X1013 vg) via the retro-orbital route. The rat pups were then returned to the mother by rubbing them with the nesting material to prevent pup rejection. Finally, the pups were sacrificed, and the livers were harvested at age P5, as most pups died at P7, to be fixed in 4% PFA and stained.

### Isolation of freshly isolated PHH for benchmarking

A fresh healthy human transplant rejected liver was harvested and cut into 1 g pieces. The liver pieces were chopped into a fine paste like consistency and submerged in Liver Digest Medium (Gibco, 17703034) for 15 min at 37°C to isolate single cells. The cells were then passed through a 100 μm strainer on ice and centrifuged at 50 × g for 3 min at 4°C. Next the H40 fraction was isolated by passing the cells through a 40 μm strainer on ice again, while the H20 fraction was isolated by passing the cells through a 20 μm strainer on ice. Finally, the isolated cells were immediately used for gene expression profiling using RT-qPCR and epigenetic profiling by ChIP.

### Organoid transplantation at the base of the liver

The HLOs and mZ-HLOs were harvested right after co-culture on Day 23 and dissociated into chunks by repeated pipetting, washed with PBS and resuspended with HCM containing 2% FBS and CEPT cocktail to increase viability. On the day of surgery, the RRG rats were fully anesthetized, and an exploratory laparotomy was performed via midline incision followed by bowel evisceration to expose the portal triad, including the portal vein. The bile duct was ligated using nylon suture proximally and distally. The HLOs and mZ-HLOs (5 × 10^6^ cells) were then transplanted orthotopically at the base of the liver in close proximity to the portal vein using TISSEEL fibrin glue (Baxter). Conversely, for Z1- and Z3-HLO affinity testing, the HLOs were injected through the portal vein or inferior vena cava using a 32G 1 inch needle in a 200 μL infusion. Bleeding was controlled by application of a bulldog clamp distal to the site of injection. The incision was then closed in two layers with 5-0 vicryl coated surgical sutures (Ethicon) and GLUture (Zoetis), and Buprenorphine (0.1 mg/kg) was administered as an analgesic. The animal was maintained on ad libitum dox chow until the day of harvest. Blood was collected regularly by the retro-orbital method as needed before the liver was harvested on Day 30.

### Immunostaining

The organoid samples were fixed with 4% paraformaldehyde for 2 hours and washed with PBS for 10 mins three times. The samples were hydrated and permeabilized with 0.1% Triton X-100/PBS and then blocked with 5% normal donkey serum. The samples were incubated with primary and secondary antibodies (as listed in Table) overnight at 4°C with gentle shaking. The samples were counterstained with DAPI, washed with PBS and cleared with refractive index matching solution (RIMS) as needed. Finally, the samples were imaged on Nikon A1R Inverted LUNV Confocal Laser Scanning Microscope.

### Protein expression assays

Albumin secretion was measured by collecting 200 μl of the supernatant from the HLOs cultured in HCM and stored at −80°C until use. The supernatant was assayed with Human Albumin ELISA Quantitation Set (Bethyl Laboratories, Inc) according to the manufacturer’s instructions. For murine GULO expression assay, the organoids were dissociated and washed with PBS. The cells were then lysed with RIPA Lysis and Extraction Buffer and Halt Protease and Phosphatase Inhibitor Cocktail (Thermo Scientific) to extract total protein and assayed with Mouse GULO/L-Gulonolactone Oxidase ELISA Kit (MyBioSource.com, MBS2890736) according to the manufacturer’s instructions.

### Metabolite assays

Bilirubin levels were measured by collecting the supernatant from HLOs treated with bilirubin and serum from the rats. The supernatant and serum were assayed with Bilirubin Assay Kit (Total and Direct, Colorimetric) (abcam, ab235627) and Bilirubin Assay Kit (Sigma-Aldrich, MAK126) according to the manufacturer’s instructions. Cellular antioxidant levels were measured by harvesting the HLOs, washing in PBS, and plating them into a 96 well assay plate. The levels were then quantitated using Cellular Antioxidant Assay Kit (ab242300) according to the manufacturer’s instructions. The nitrogen related metabolite assays were carried out by harvesting the HLOs, washing in PBS, and plating them into a 96 well assay plate. The Glutathione, Ammonia, Urea, Glutamine, Glucose, and Triglyceride levels were then assayed by using the corresponding Glutathione, Ammonia, Urea, Glutamine, Glucose, and Triglyceride assay kits (abcam ab65322, ab83360, ab83362, ab197011, ab65333, ab65336) according to the manufacturer’s instructions.

### Metabolic activity assays

CYP3A4 and CYP1A2 assays were performed by harvesting the HLOs, washing in PBS, plating them into a 96 well assay plate, and treating them with rifampicin and omeprazole respectively for 24 hrs. The assays were then performed using P450-Glo CYP3A4 and CYP1A2 Assay (Promega, V8802 and V8422) and normalized using CellTiter-Glo Luminescent Cell Viability Assay (G7572) according to the manufacturer’s instructions. The Notch1 assay was carried out by transfecting the HLOs with the experimental, reporter and negative vectors from the Human Notch1 Pathway Reporter kit (amsbio, 79503) using Lipofectamine 3000 and Opti-MEM I according to the manufacturer’s instructions. Dual Luciferase Assay System (amsbio, 60683- 2) for Notch1 assay was then used to measure the Firefly luciferase activity and compared to Renilla luciferase activity to normalize the transfection efficiency. The luciferase assay indicates Notch activity using a CSL (CBF1/ RBPJK) luciferase reporter vector, Notch pathway responsive reporter. Notch1 is cleaved by gamma secretase and NICD is released into the nucleus which is detected by the luciferase reporter as active Notch signaling. The nitrogen metabolism related enzyme assays were carried out by harvesting the HLOs, washing in PBS, and plating them into a 96 well assay plate. The GS activity and GST activity levels were then assayed by using the Glutamine Synthetase Activity, Glutathione S Transferase Activity, Lipase Activity, and Glucokinase Activity Assay Kit (abcam ab284572, ab65325, ab102524, ab273303) according to the manufacturer’s instructions. The apoptosis assay was carried out by lysing the HLOs and assaying the lysate with a Caspase-3 Assay Kit (Colorimetric) (ab39401) according to the manufacturer’s instructions. Finally, rat serum was assayed with AST and ALT Activity Assay Kit (Sigma-Aldrich, MAK055 and MAK052) and quantified by a BioTek^®^ Synergy H1 plate reader.

### Zonal toxicity assay

The HLOs were induced with 3-MC (50 μM) for alcohol degradation and drug conjugation metabolism 24 hours prior to the toxicity assay. After induction a toxic dose of the zone 1 toxin allyl alcohol (200μM) was supplemented for 2hr at 37°C. On the other hand, a toxic dose of the zone 3 toxin acetaminophen (10mM) was supplemented for 4hr at 37°C on different batches. However, for the mZ-HLO regenerative potential assay the toxins were incubated for 1 hr at 37°C. Subsequently, organoids were supplied with fresh media and the organoids were fixed after 24 hr in 4% PFA and stained. The cultures were then tested for Caspase 3 activity using the cellular lysate collected from the organoid culture. Separately, the cultures were also tested for viability using CellTiter-Glo Luminescent Cell Viability Assay.

### Quantification and statistical analysis

Statistical analyses were mainly performed using R software v4.2.0 with unpaired two-tailed Student’s t-test, one-way Anova and post hoc Tukey’s test, or Welch’s t-test. Statistical analyses for non-normally distributed measurements were performed using non-parametric Kruskal-Wallis and post hoc Dunn-Holland-Wolfe test. For comparisons between unpaired groups, when groups were independent and the variances were unequal, non-parametric Brunner-Munzel test was performed, unless noted otherwise. P values < 0.05 were considered statistically significant. n value refers to biologically independent replicates. The image analyses were non-blinded. Statistical parameters are found in the figures and Figure legends where ns = P > 0.05, * = P ≤ 0.05, ** = P ≤ 0.01, *** = P ≤ 0.001, and **** = P ≤ 0.0001. Using G*Power software, for each experiment, we determined the minimum sample size to collect the data for using the preliminary effect sizes, α = 0.05 and power = 0.8. For each experimental data, we also conducted a post hoc power analysis to determine whether our design had enough power. We had enough power (power > 0.8) for all our experiments.

### Data and code availability

The RNA-seq, ChIP-seq and snRNA-seq data reported in this paper have been deposited to NCBI Gene Expression Omnibus (GEO) with the following accession number: GSE222654. Reference codes for bioinformatics analyses can be found at the following link: https://github.com/hasanwraeth. All other data supporting the findings of this study are available from the corresponding author on reasonable request.

## Acknowledgements

We appreciate Asuka Kodaka for scientific illustration materials used in this study. We would like to thank all Takebe, Zorn, Helmrath and Wells lab members for their support and feedback. We would also like to acknowledge the CCHMC Confocal Imaging Core (RRID:SCR_022628), Pathology Research Core (RRID:SCR_022637), Pluripotent Stem Cell Facility (RRID:SCR_022634), Transgenic Animal and Genome Editing Core (RRID:SCR_022642), Veterinary Services Facility, and Single Cell Genomics Core (RRID:SCR_022653) for providing gene expression profiling and analysis and Andrew Potter for expertise in single cell isolation and troubleshooting. Finally, we would also like to acknowledge the thesis committee (Jorge A. Bezerra, MD; Vivian Hwa, PhD; Ralph Cutler C. Quillin, MD; Jason Tchieu, PhD; James M. Wells, PhD) for HAR for their continued support and constructive criticism.

## Author contributions

H.A.R. designed and performed research, analyzed data, performed the bioinformatics analyses, and wrote the paper. C.S., A.A.R., S.S., K.G. conducted research and revised the paper. K.I., A.B., and J.M. designed research and revised the paper. T.T. designed research and wrote the paper.

## EXTENDED DATA FIGURE LEGENDS

**Extended Data Fig. 1.**
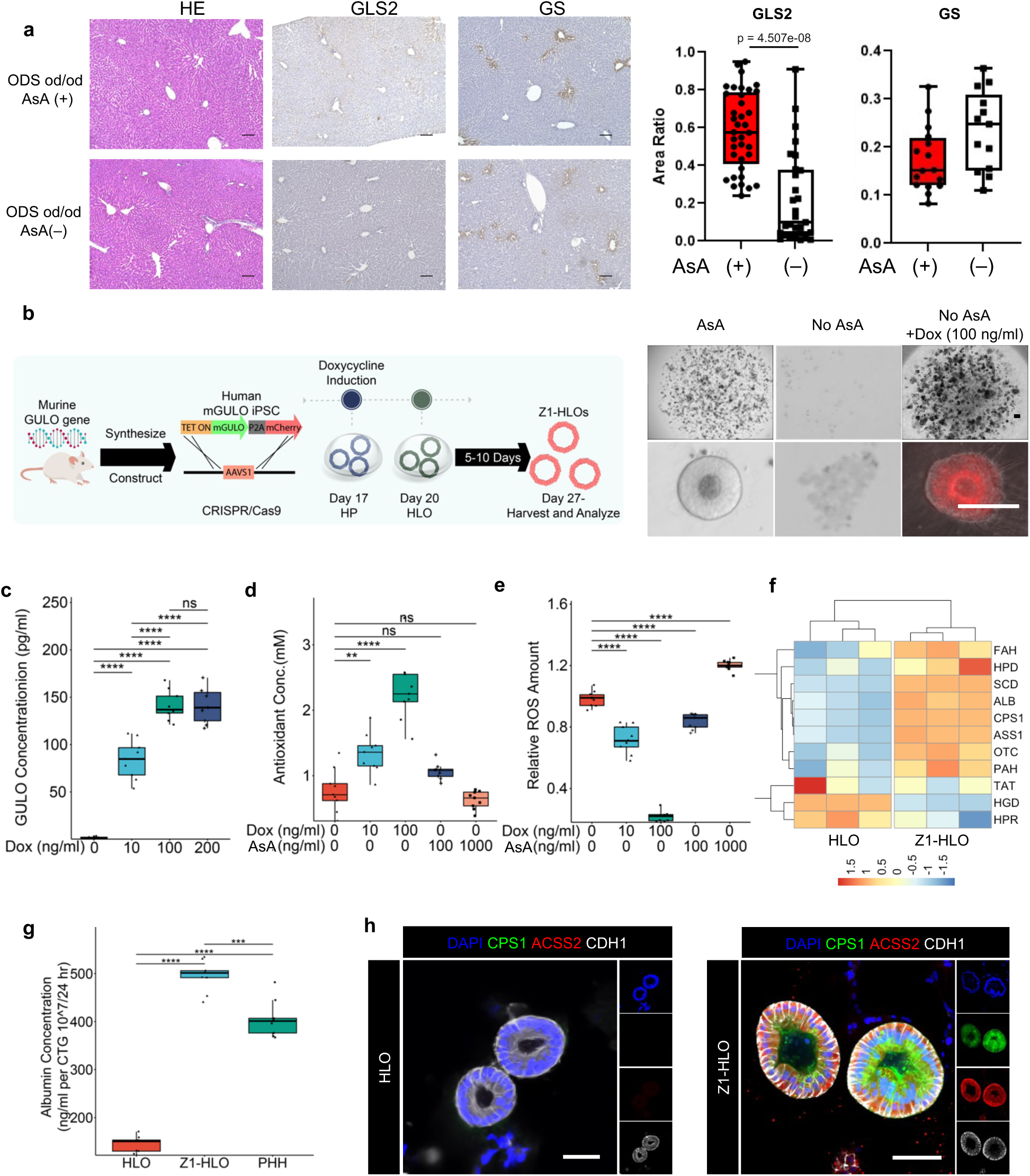
Intracellular redox management enables CPS1+ hepatocyte specification in HLOs. a) H&E histology, GLS2 IHC, and GS IHC images of liver sections are shown in panels (ODS od/od (GULO mutant) rat treated with 0.2% Ascorbic acid (AsA), ODS od/od rat treated without AsA. Scale bars indicate 100µm. The graph shows the GLS2 or GS positive area ratio versus the hematoxylin positive area in ODS od/od (GULO mutant) rat treated with 0.2% Ascorbic acid (AsA) and ODS od/od rat treated without AsA (n=2 ; Mean +-SD, Mann-Whitney *U* test, p = 4.507e-08) Data points are shown GLS2 area at portal vein (37 portal vein sections of + AsA rats, 27 portal vein sections of - AsA rats) and GS area in 8 images of +/- AsA rat for analysis with Fiji. b) Schematic for development of Z1-HLOs and doxycycline induction to induce CPS1+ hepatocyte specification (left). Brightfield and fluorescence images of mCherry expression in ascorbate depleted Dox (100 ng/ml) treated Z1-HLOs compared to HLOs with ascorbic acid depletion at day 20 and control HLOs (right). c) ELISA for mGULO protein concentration in Dox treated Z1-HLOs compared to control HLOs. (n = 9 independent experiments) d) Cellular Antioxidant concentration in Dox treated Z1-HLOs compared to control HLOs. (n = 9 independent experiments) e) ROS levels in Dox treated and extracellular ascorbate induced Z1-HLOs compared to control HLOs. (n = 9 independent experiments) f) Heatmap of Zone 1 genes from RNAseq dataset for Dox treated Z1-HLOs compared to control. g) Albumin ELISA for Z1-HLOs treated with Dox compared to control HLOs and PHH normalized by cell viability. (n = 9 independent experiments) h) Immunofluorescence images of Dox treated Z1-HLOs for CPS1, ACSS2 and CDH1 compared to control HLOs. Scale bar indicates 200 µm.

**Extended Data Fig. 2.**
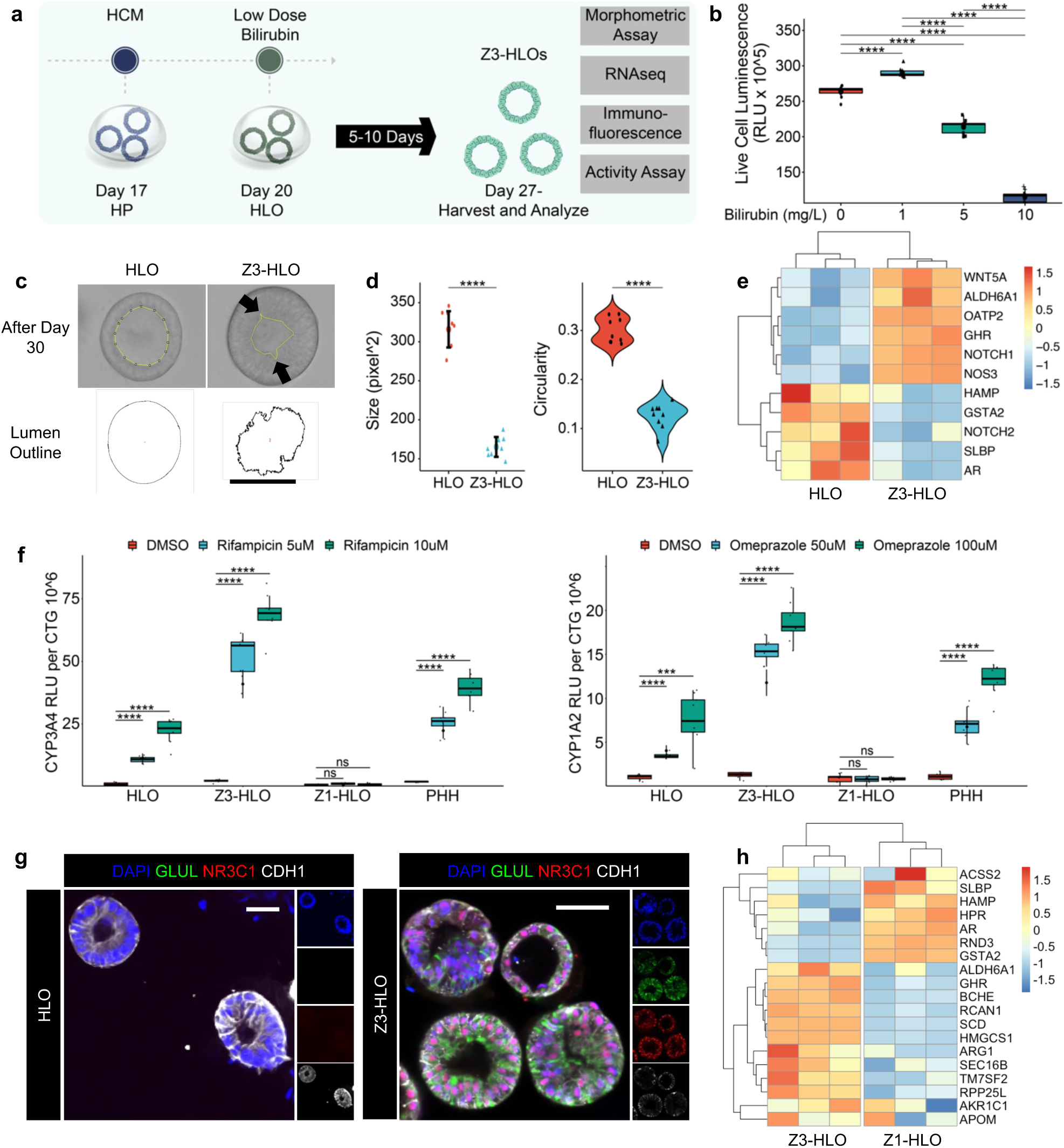
Low dose bilirubin promotes GLUL+ hepatocyte specification in HLOs. a) Schematic for low dose bilirubin treatment and Z3-HLO development to induce GLUL+ expression. b) Cell viability assay with different concentration of bilirubin to titrate dose for maximal viability. (n = 9 independent experiments). c) Brightfield image of Z3-HLOs treated with low dose bilirubin (1mg/L) compared to control, and luminal outline using ImageJ, arrows indicate luminal projections that are similar to bile canaliculi found in human liver. Scale bar indicates 200 µm. d) Comparison of size and circularity of lumen of the control and 1 mg/L bilirubin treated Z3-HLOs. (n = 9 independent experiments) e) Heatmap of Zone 3 genes from RNAseq dataset for bilirubin treated Z3-HLOs compared to control. f) CYP3A4 activity assay in response to Rifampicin in control, Z1-, Z3-HLOs, and PHH (left). CYP1A2 activity assay in response to Omeprazole in control, Z1-, Z3-HLOs, and PHH (right). (n = 9 independent experiments). g) Immunofluorescence images of Z3-HLOs for GLUL, NR3C1 and CDH1 compared to control HLOs. Scale bar indicates 200 µm. h) Heatmap of Z1- and Z3- HLOs depicting expression of zonal genes and lack of consensus expression of markers such as *ARG1* and *AKR1C1*.

**Extended Data Fig. 3.**
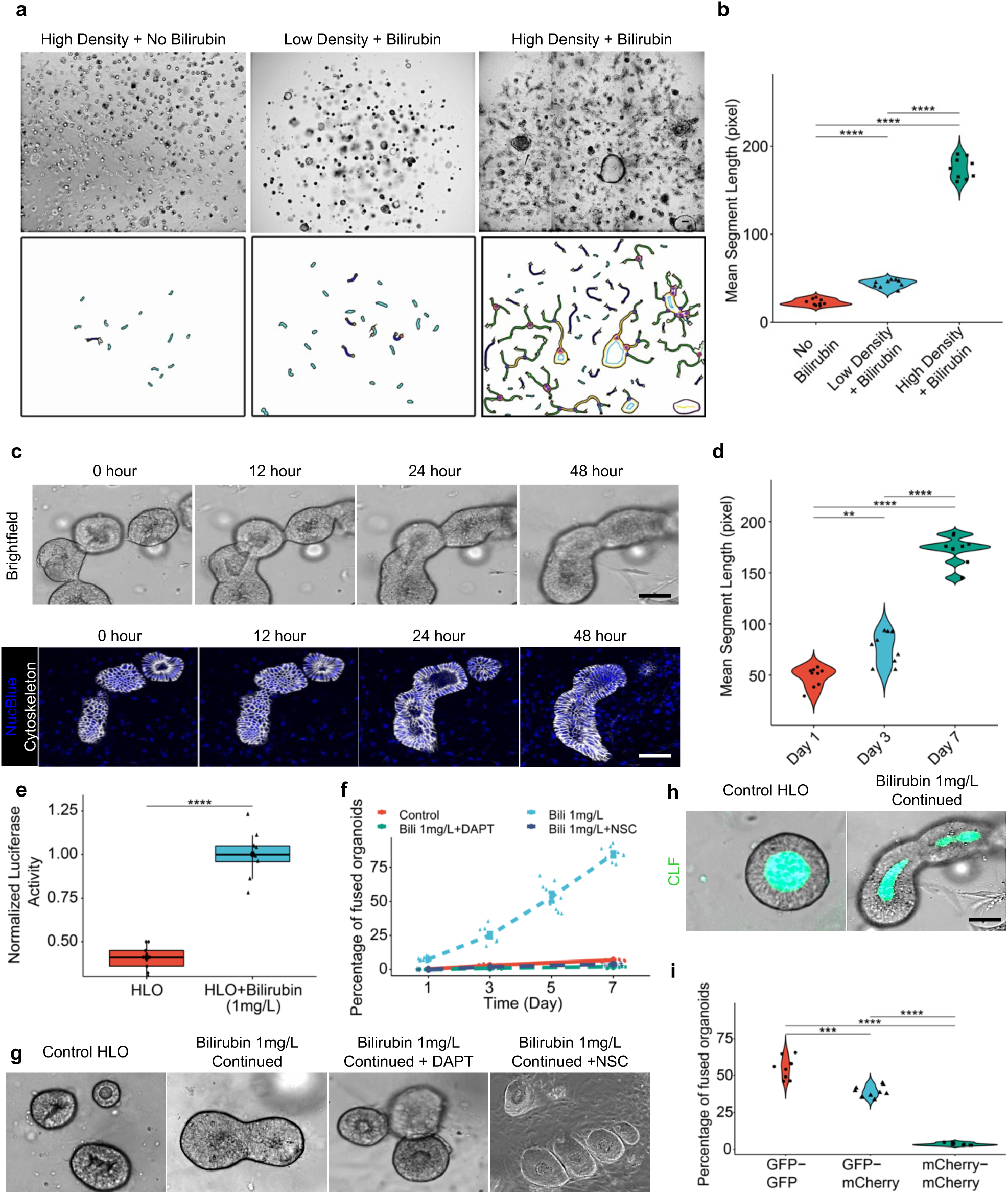
Bilirubin induced fusion requires close proximity and cytoskeletal signaling. a) Brightfield images of bilirubin induced fusion in high density HLOs compared to low density and no bilirubin treatment. Scale bar indicates 200 µm. b) Comparison of mean segment length in high density HLOs compared to low density and no bilirubin treatment. (n = 9 independent experiments) c) Brightfield and live staining images (NucBlue: Blue, Cytoskeleton: White) show progression of organoid fusion after continued treatment with bilirubin (1 mg/L). Scale bar indicates 200 µm. d) Comparison of mean segment length of the HLOs from Day 1 to Day 7. (n = 9 independent experiments). e) NOTCH activity assay in bilirubin treated HLOs compared to control. (n = 9 independent experiments). f) Percentage of fused organoids after bilirubin treatment in DAPT (Notch inhibitor) and NSC (Ezrin Inhibitor, NSC668394) treated HLOs compared to control. (n = 9 independent experiments). g) Brightfield images of bilirubin induced fused HLOs compared to DAPT or NSC668394 treatment and control HLOs. Scale bar indicates 200 µm. h) CLF assay for self-assembled organoids compared to control. Scale indicates 200 µm. i) Percentage of fused organoids for each type of organoid. (n = 9 independent experiments).

**Extended Data Fig. 4.**
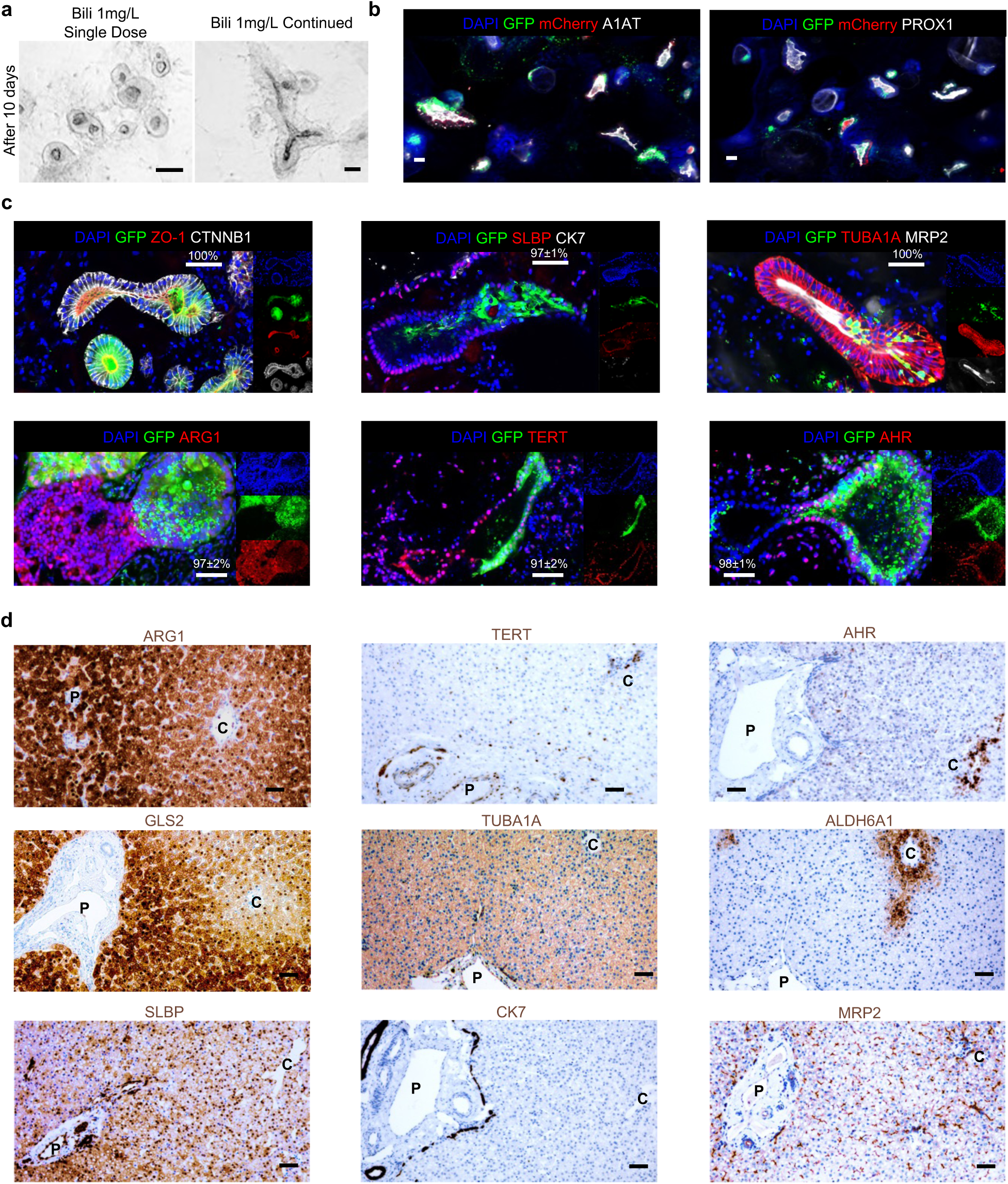
Immunostaining of mZ-HLOs compared to neonatal liver show similar features. a) Brightfield images of fused HLOs following continued treatment with bilirubin compared to single dose treatment after 10 days. b) Immunofluorescence images (bottom) of mZ-HLOs depicting GFP, mCherry, PROX1 and A1AT. Scale bar indicates 200 µm. c) Immunofluorescence images of mZ-HLOs for pan liver markers: TUBA1A, CTNBB1, luminal marker: ZO-1 (depicting continuous lumen); Zone 1 markers: ARG1 and SLBP; Zone 2 marker TERT; Zone 3 markers: AHR, and MRP2; and Cholangiocyte marker CK7. Scale bar indicates 200 µm. Numbers indicate the ratio of correctly fused organoids and total number of organoids. d) Immunohistochemistry images of neonatal liver sections for pan liver marker TUBA1A; Cholangiocyte marker CK7; Zone 1 markers: ARG1, SLBP, and GLS2; Zone 2 marker TERT; and Zone 3 markers: AHR, ALDH6A1 and MRP2. Scale bar indicates 200 µm. (P: Portal vein, C: Central vein).

**Extended Data Fig. 5.**
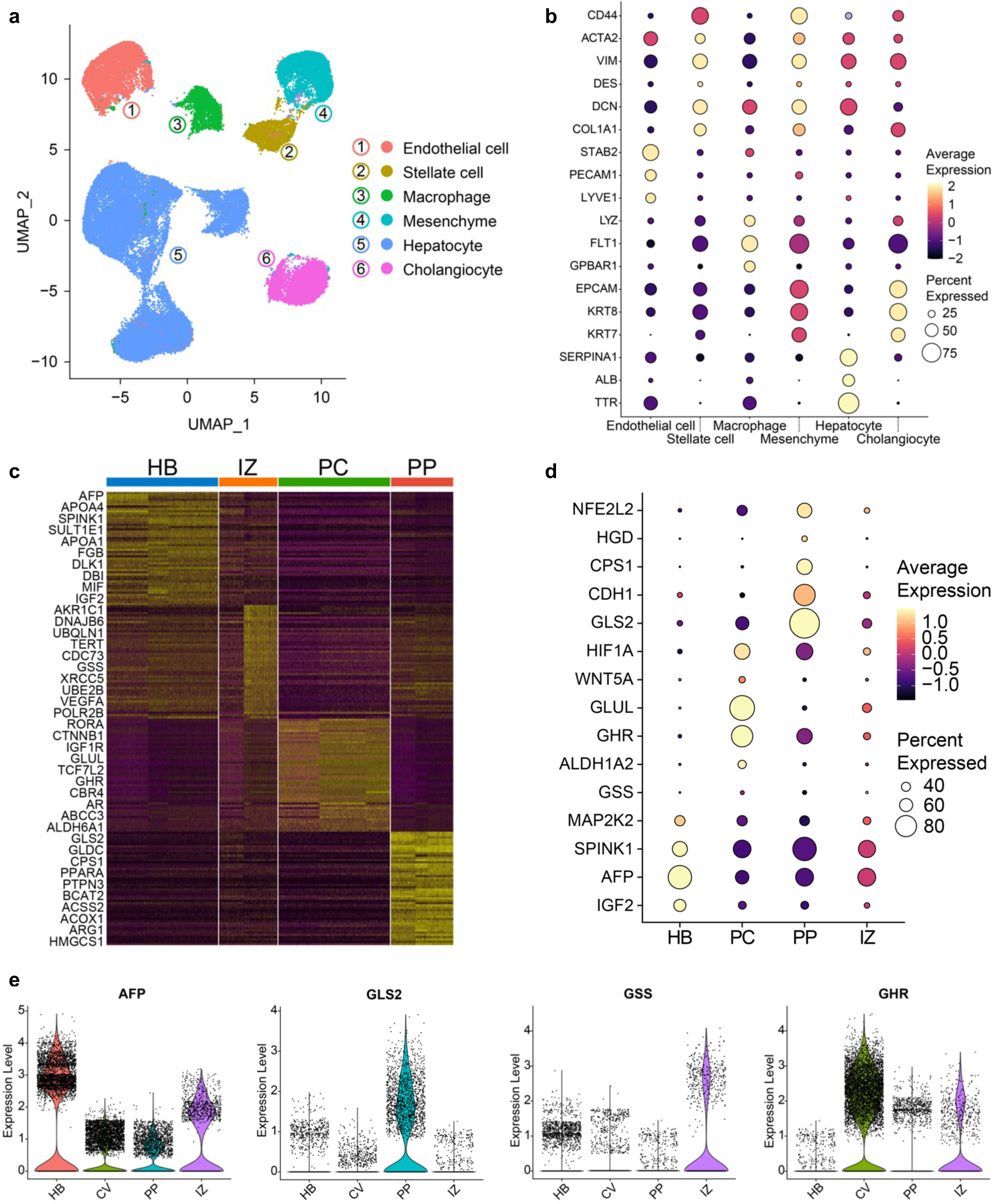
Single cell profiling of mZ-HLOs indicate the emergence of zonal like populations. a) UMAP plot with the major populations (hepatocytes, cholangiocytes, endothelial cells, macrophages, stellate cells, and mesenchyme) of all nuclei in mZ-HLOs. b) Distinct expression profile all populations in mZ-HLOs. The size of the circle indicates the percentage of nuclei in each population expressing each gene. The color represents the average expression level for the indicated gene. c) Heatmap showing scaled mean expression of all genes in each cluster. Top 10 marker genes in each cluster have been added as labels. d) Expression of known hepatoblast and zonal hepatocyte marker genes in each population. e) Violin plot for expression of AFP (hepatoblast gene), GSS (interzonal hepatocytes), GHR (pericentral hepatocyte), and GLS2 (periportal hepatocyte).

**Extended Data Fig. 6.**
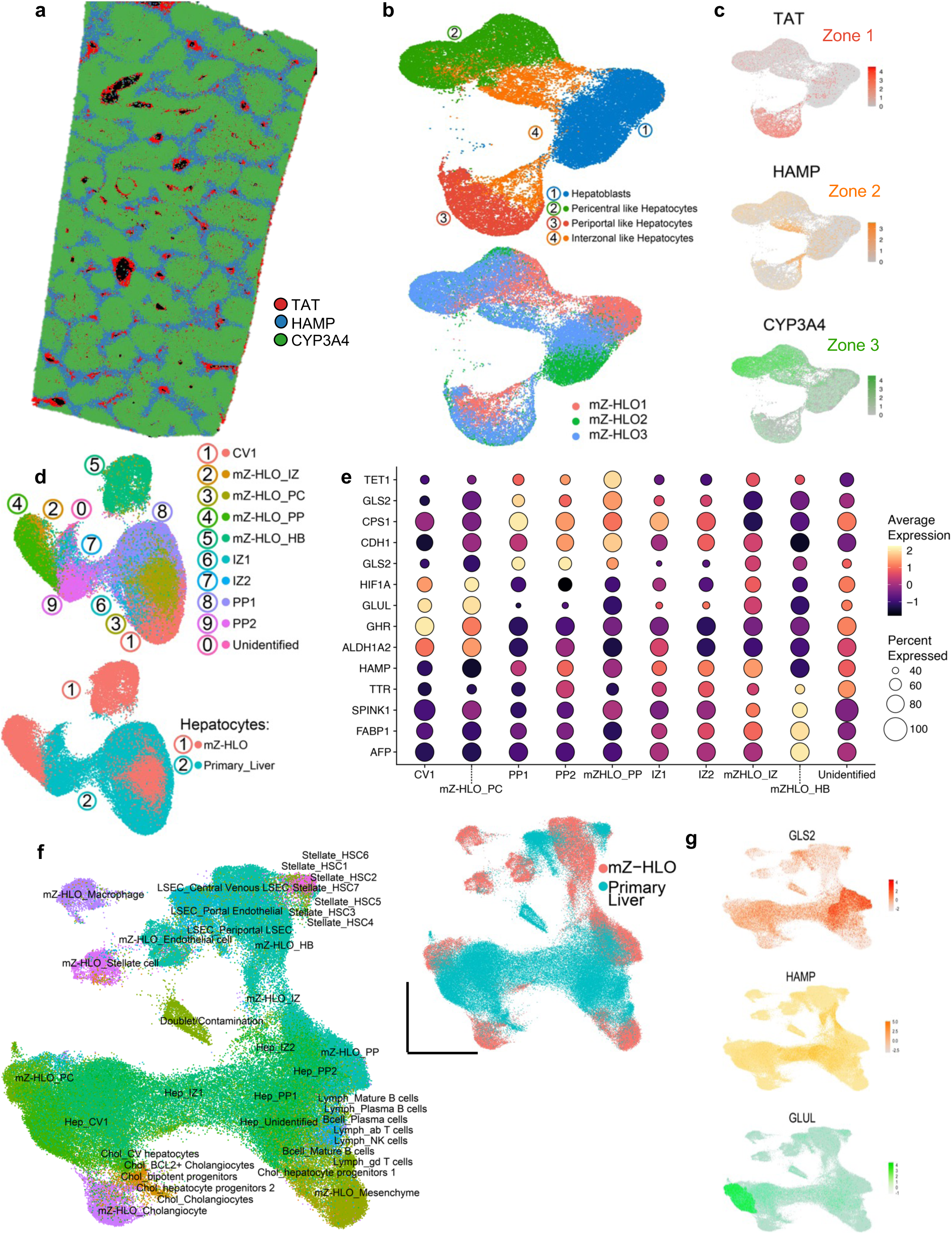
Pseudo-spatial profiling of mZ-HLOs show similarity of zonal expression to primary liver tissue. a) Spatial plot for *TAT* (zone 1), *HAMP* (zone 2), and *CYP3A4* (zone 3) markers in 10X Xenium healthy human liver dataset (publicly available dataset). b) UMAP plot of mZ-HLO with hepatocyte populations (top) and distribution of replicate data (bottom). c) Feature plot for *TAT* (zone 1), *HAMP* (zone 2), and *CYP3A4* (zone 3) markers. d) UMAP plot for zonal hepatocyte populations from primary liver (Andrews *et al.*, 2022) and mZ-HLOs integrated together (top). UMAP plot depicting distribution for total hepatocyte populations from primary liver and mZ-HLOs integrated together (bottom). e) Expression of known hepatoblast and zonal hepatocyte marker genes in mZ-HLOs benchmarked against Andrews *et al.*, 2022 snRNAseq dataset. f) UMAP plot for all cell types (inset: sample distribution) from primary liver datasets and mZ-HLOs integrated together. g) Feature plot for *GLS2* (zone 1), *HAMP* (zone 2), and GLUL (zone 3) markers.

**Extended Data Fig. 7.**
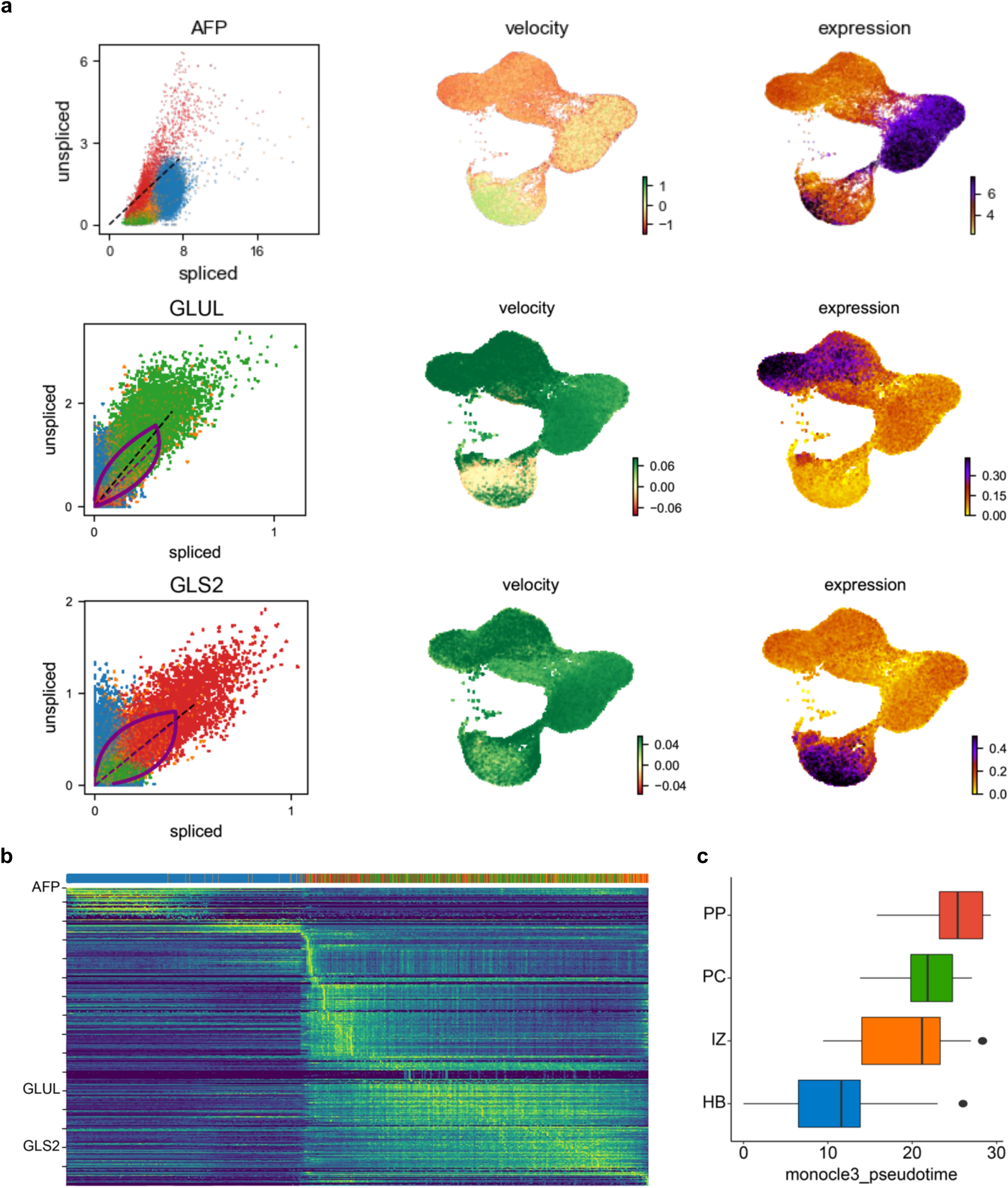
RNA velocity and pseudotime analysis in mZ-HLOs. a) Phase portrait of *AFP, GLUL, and GLS2* depicting the dynamics of the gene splicing in the nuclei with the velocity and expression of *AFP, GLUL, and GLS2* in nuclei as feature plots. b) SOM (Self Organizing Map) of single-nuclei transcriptome-derived zonation profiles for mZ-HLOs based on the different populations. c) Boxplot showing the pseudotime of each nuclei population in mZ-HLOs.

**Extended Data Fig. 8.**
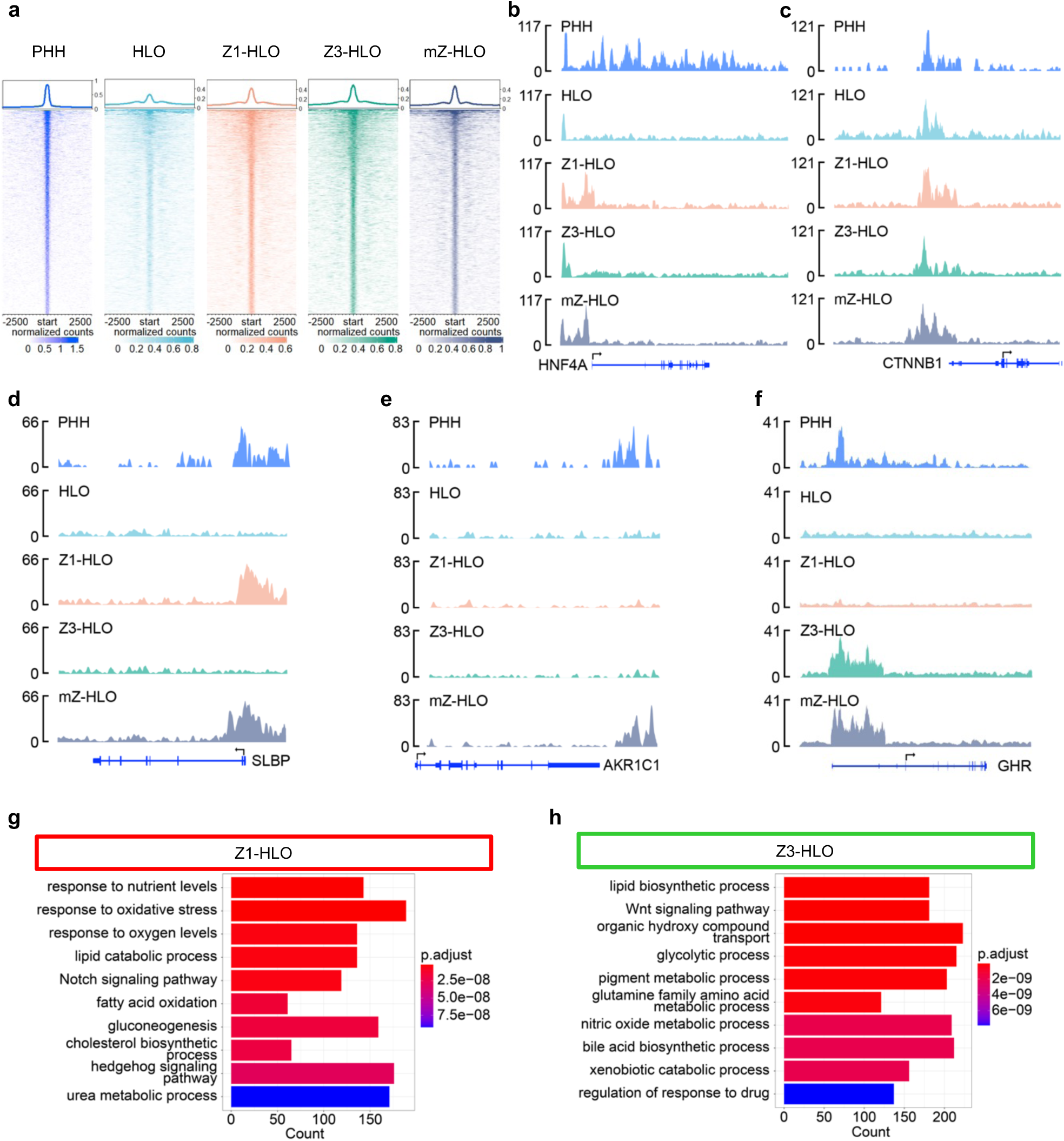
EP300 differentially regulates zonal genes in mZ-HLOs in conjunction to distinct transcription factors. a) Peak density plots showing EP300 bound loci, a marker of active enhancers. Profile plot of all peaks are in the top panel. b) Genome browser view of HNF4A (pan marker) showing the EP300 ChIPseq peak. c) Genome browser view of CTNNB1 (pan marker) showing the EP300 ChIPseq peak. d) Genome browser view of SLBP (zone 1 gene) showing the EP300 ChIPseq peak. e) Genome browser view of AKR1C1 (zone 2 gene) showing the EP300 ChIPseq peak. f) Genome browser view of GHR (zone 3 gene) showing the EP300 ChIPseq peak. g) Top 10 upregulated Gene Ontology terms (Biological Process) for the gene regulated bound by EP300 in the Z1-HLOs. h) Top 10 upregulated Gene Ontology terms (Biological Process) for the gene regulated bound by EP300 in the Z3-HLOs.

**Extended Data Fig. 9.**
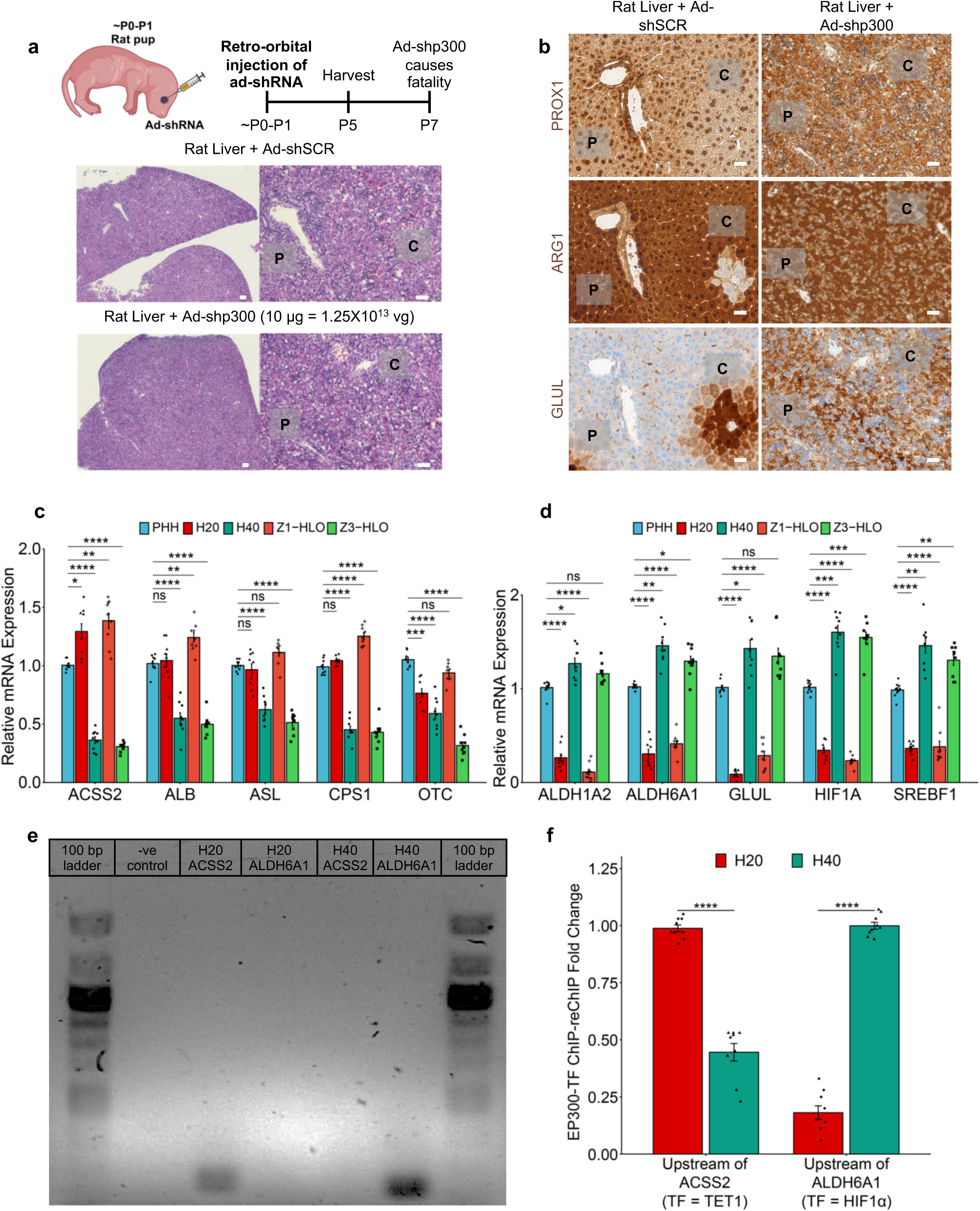
EP300 and partner transcription factors are important for zonal liver development. a) Experimental timeline for testing role of EP300 in zonal liver development using Ad-shp300 (adenoviral vector for p300 shRNA, top). H&E stain for sections from rat liver injected with Ad-shSCR (adenoviral vector for scrambled shRNA, middle) and Ad-shp300 (adenoviral vector for p300 shRNA, bottom). Scale bar indicates 200 µm. (P: Portal vein, C: Central vein) b) ICH stain for PROX1, ARG1, and GLUL of sections from rat liver injected with Ad-shSCR (adenoviral vector for scrambled shRNA, top) and Ad-shp300 (adenoviral vector for p300 shRNA, bottom). Scale bar indicates 200 µm. (P: Portal vein, C: Central vein) c) RT-qPCR of *ALB, ACSS2, ASL, CPS1,* and *OTC* (zone 1) gene for Z1- and Z3-HLOs compared to freshly isolated PHH, H20 (20 um periportal hepatocytes) and H40 (40 um pericentral hepatocytes) (Data is mean ± SD, n = 9 independent experiments). d) RT-qPCR of *ALDH1A2, ALDH6A1, HIF1A, SREBF1,* and *GLUL* (zone 3) gene for Z1- and Z3-HLOs compared to freshly isolated PHH, H20 (20 um periportal hepatocytes) and H40 (40 um pericentral hepatocytes) (Data is mean ± SD, n = 9 independent experiments). e) EP300-TF ChIP-reChIP-PCR for H20 (20 um periportal hepatocytes) and H40 (40 um pericentral hepatocytes). f) NR3C1-MECP2 ChIP-reChIP-qPCR for samples in (e). Data are mean ± SD, n = 9 independent experiments.

**Extended Data Fig. 10.**
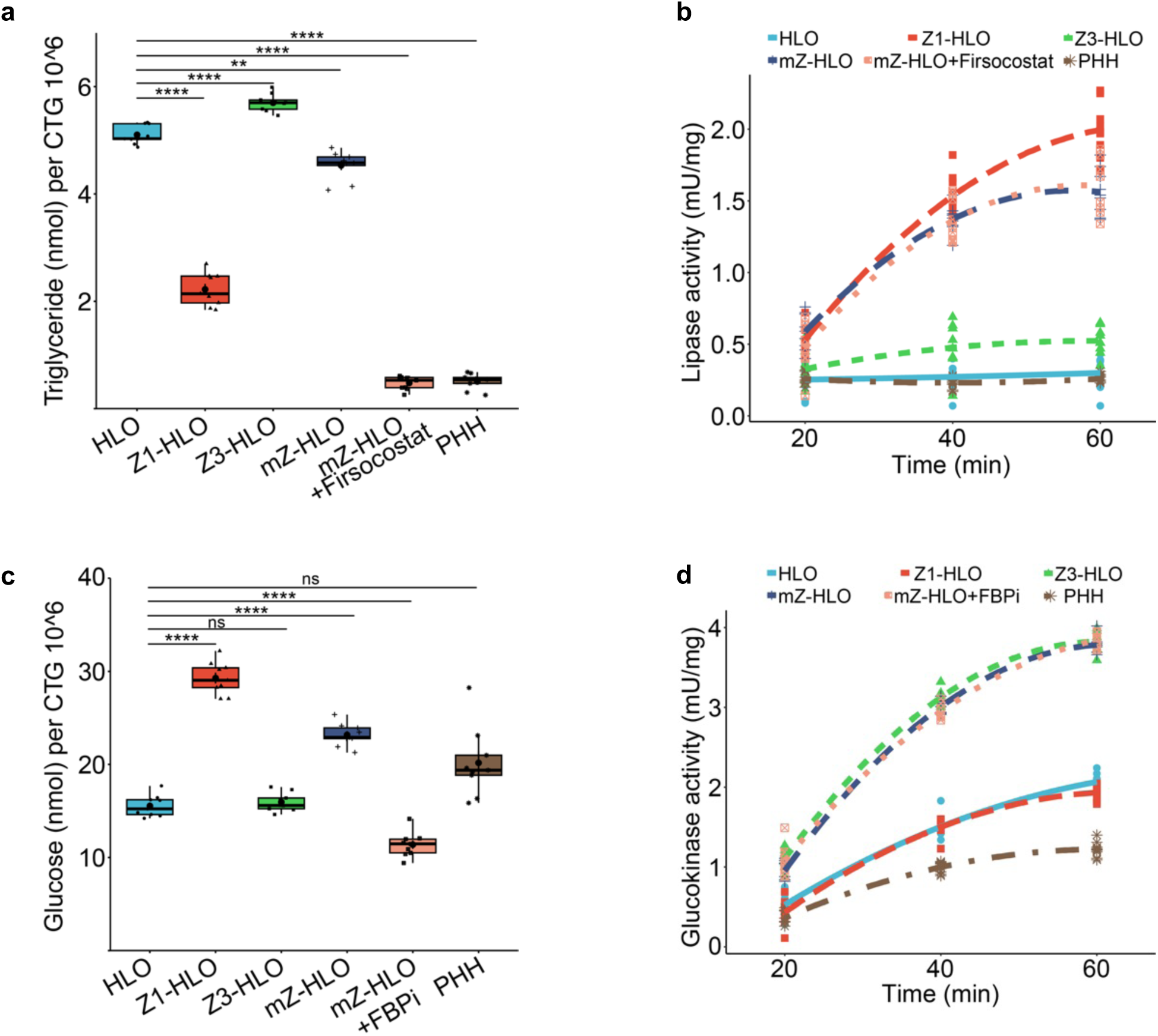
Interzonal dependent lipid and glucose metabolism in mZ-HLOs. a) Triglyceride assay for mZ-HLOs with and without Firsocostat treatment compared to Z1-, Z3-, control HLOs, and PHH (n = 9 independent experiments). b) Lipase activity assay for mZ-HLOs with and without Firsocostat treatment compared to Z1-, Z3-, control HLOs, and PHH (n = 9 independent experiments). c) Glucose assay for mZ-HLOs with and without FBPi treatment compared to Z1-, Z3-, control HLOs, and PHH (n = 9 independent experiments). d) Glucokinase activity assay for mZ-HLOs with and without FBPi treatment compared to Z1-, Z3-, control HLOs, and PHH (n = 9 independent experiments).

**Extended Data Fig. 11.**
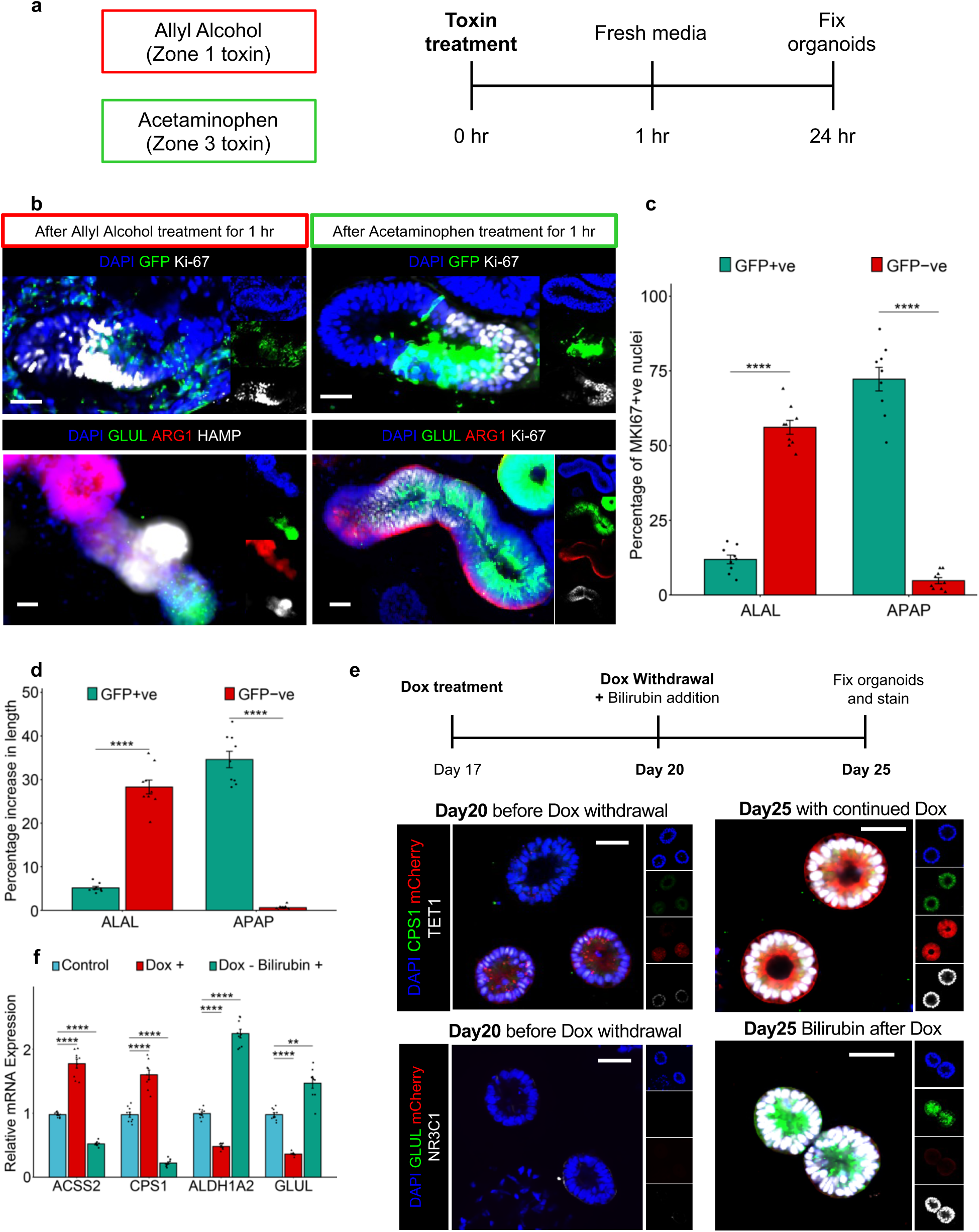
mZ-HLOs exhibit zone specific regenerative potential in response to toxins. a) Experimental timeline for testing zonal regenerative potential of mZ-HLOs in response to zone specific toxins. b) Immunofluorescence images of mZ-HLOs for proliferative marker Ki-67, GLUL, ARG1 and HAMP in response to Allyl Alcohol (Zone 1 toxin, left). Immunofluorescence images of mZ-HLOs for proliferative marker Ki-67, GLUL and ARG1 in response to Acetaminophen (Zone 3 toxin). Scale bar indicates 200 µm. c) Comparison of Ki-67 + nuclei in different fluorescent regions in response to zone specific toxins. (n = 9 independent experiments) d) Comparison of length of different fluorescent regions in response to zone specific toxins. (n = 9 independent experiments) e) Immunofluorescence images of CPS1, TET1, GLUL, NR3C1, and mCherry in Z1-HLOs with Dox treatment and after Dox withdrawal and persistent bilirubin treatment at Day 20 and 25. Scale bar indicates 200 µm. f) RT-qPCR of *ACSS2, CPS1, ALDH1A2* and *GLUL* gene for Z3-HLOs with Dox (Dox +) and with bilirubin after Dox withdrawal (Dox – Bilirubin +) compared to control HLOs (Data is mean ± SD, n = 9 independent experiments).

**Extended Data Fig. 12.**
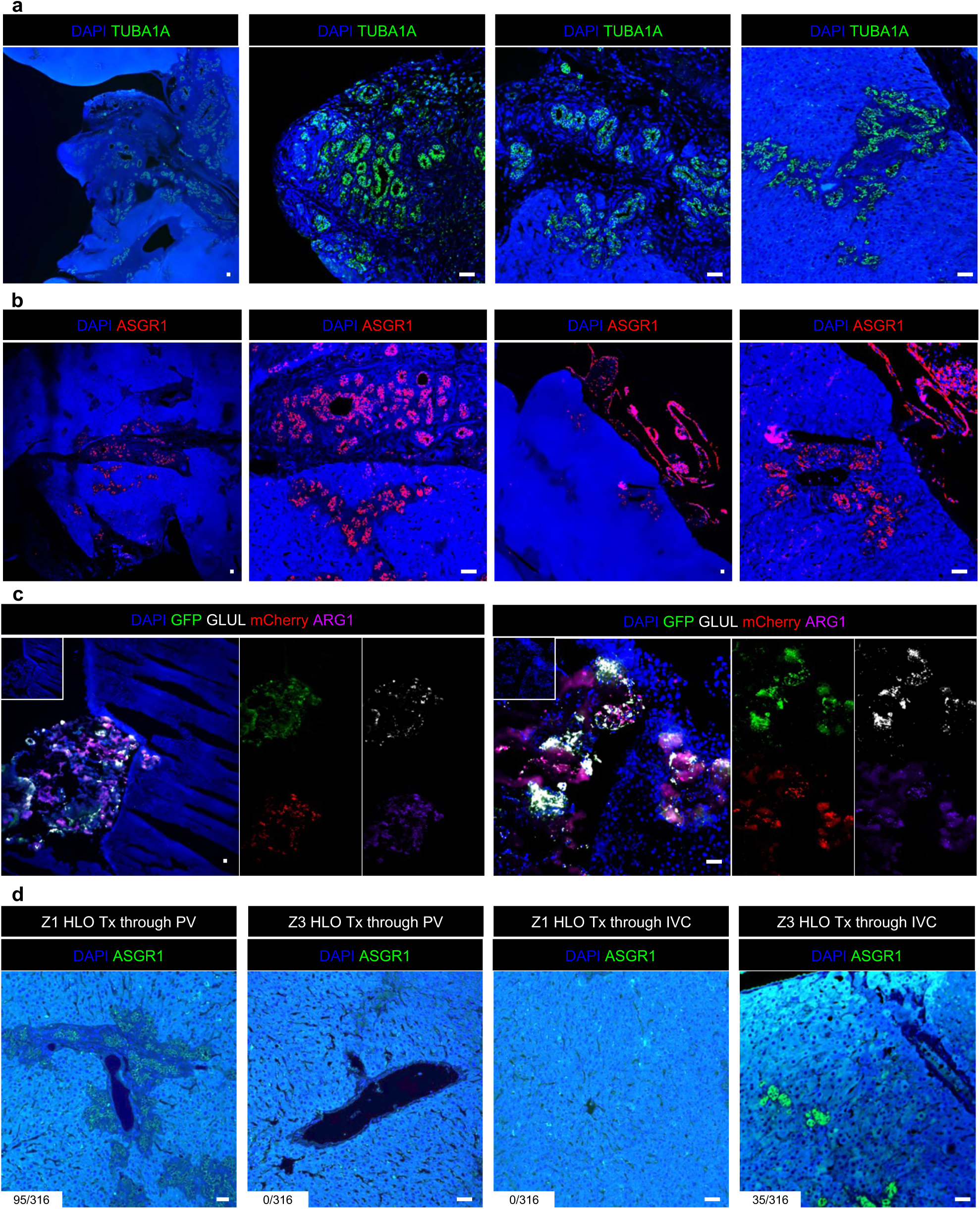
mZ-HLOs invade into the liver parenchyma of RRG rats after transplantation. a) Immunofluorescence images for human TUBA1A of mZ-HLOs transplanted in RRG rat liver. Scale bar indicates 200 µm. b) Immunofluorescence images for human ASGR1 of mZ-HLOs transplanted in RRG rat liver. Scale bar indicates 200 µm. c) Immunofluorescence images for GFP, mCherry, human ARG1 and GLUL of mZ-HLOs transplanted in RRG rat liver. Scale bar indicates 200 µm. d) Immunofluorescence images for human ASGR1 of Z1 and Z3-HLOs transplanted in RRG rat liver through the portal vein and inferior vena cava. Scale bar indicates 200 µm. Numbers indicate the ratio of the area of integrated organoids and total area of the liver parenchyma in view in 10^3^ pixel^2^.

## Extended Data Methods Animal experiments

Three or four-week-old male ODS (ODS/Shi Jcl-od/od) rats were purchased from CLEA Japan, Inc.(Tokyo, Japan). They were housed in individual cages and maintained at temperature and humidity with 12 hours of light exposure each day from 7 a.m. to 7p.m. They were given free access to water and a purified diet. The compositions of the diet (AsA 0mg/kg, AsA-free diet) and with or without 2% AsA (Wako-Fujifilm, Japan) contained water. After 1 week or 2week of feeding, they were anesthetized with isoflurane and sampling liver with a perfusion fix of 4% Paraformaldehyde (Nacalai, Japan). Animal care and experimental procedures were approved by the Animal Research Committee of TMDU (approval number A2023554).

---

**Table S1:**
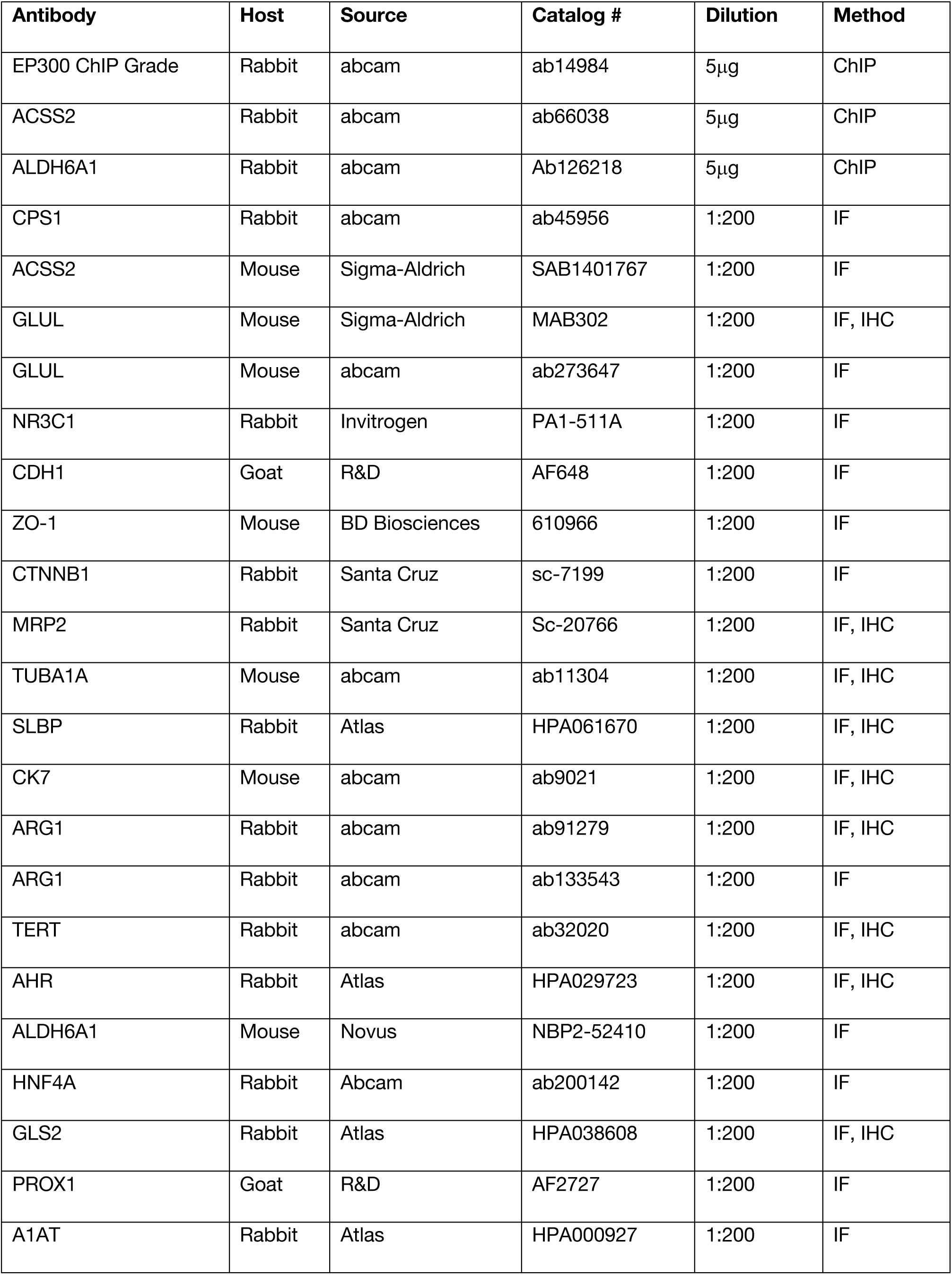
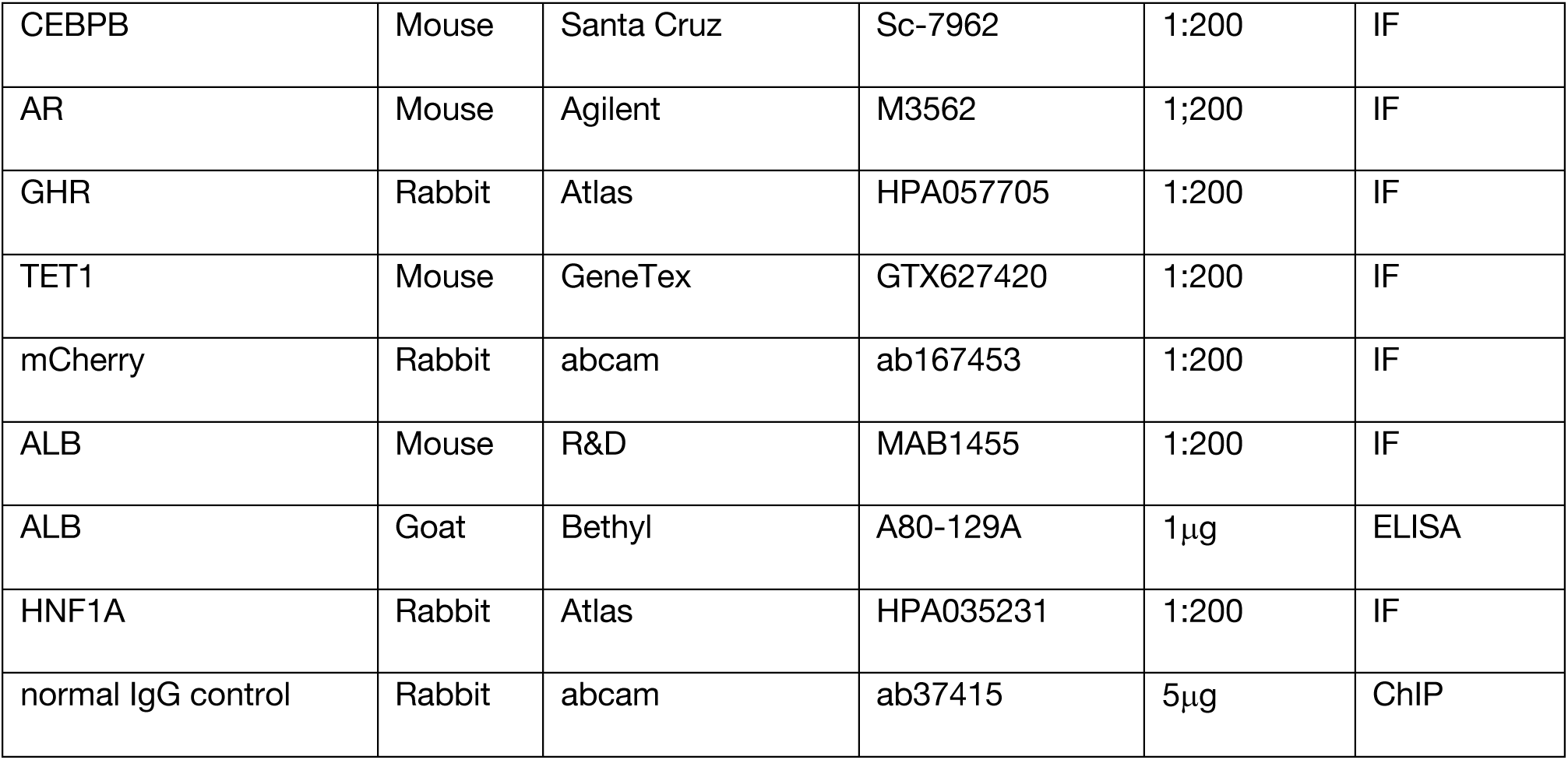
List of antibodies used for immunostaining (IC), ELISA, and ChIP-seq (ChIP) in organoid experiment.

| Antibody | Host | Source | Catalog # | Dilution | Method |
| --- | --- | --- | --- | --- | --- |
| EP300 ChIP Grade | Rabbit | abcam | ab14984 | 5µg | ChIP |
| ACSS2 | Rabbit | abcam | ab66038 | 5µg | ChIP |
| ALDH6A1 | Rabbit | abcam | Ab126218 | 5µg | ChIP |
| CPS1 | Rabbit | abcam | ab45956 | 1:200 | IF |
| ACSS2 | Mouse | Sigma-Aldrich | SAB1401767 | 1:200 | IF |
| GLUL | Mouse | Sigma-Aldrich | MAB302 | 1:200 | IF, IHC |
| GLUL | Mouse | abcam | ab273647 | 1:200 | IF |
| NR3C1 | Rabbit | Invitrogen | PA1-511A | 1:200 | IF |
| CDH1 | Goat | R&D | AF648 | 1:200 | IF |
| ZO-1 | Mouse | BD Biosciences | 610966 | 1:200 | IF |
| CTNNB1 | Rabbit | Santa Cruz | sc-7199 | 1:200 | IF |
| MRP2 | Rabbit | Santa Cruz | Sc-20766 | 1:200 | IF, IHC |
| TUBA1A | Mouse | abcam | ab11304 | 1:200 | IF, IHC |
| SLBP | Rabbit | Atlas | HPA061670 | 1:200 | IF, IHC |
| CK7 | Mouse | abcam | ab9021 | 1:200 | IF, IHC |
| ARG1 | Rabbit | abcam | ab91279 | 1:200 | IF, IHC |
| ARG1 | Rabbit | abcam | ab133543 | 1:200 | IF |
| TERT | Rabbit | abcam | ab32020 | 1:200 | IF, IHC |
| AHR | Rabbit | Atlas | HPA029723 | 1:200 | IF, IHC |
| ALDH6A1 | Mouse | Novus | NBP2-52410 | 1:200 | IF |
| HNF4A | Rabbit | Abcam | ab200142 | 1:200 | IF |
| GLS2 | Rabbit | Atlas | HPA038608 | 1:200 | IF, IHC |
| PROX1 | Goat | R&D | AF2727 | 1:200 | IF |
| A1AT | Rabbit | Atlas | HPA000927 | 1:200 | IF |
| CEBPB | Mouse | Santa Cruz | Sc-7962 | 1:200 | IF |
| AR | Mouse | Agilent | M3562 | 1;200 | IF |
| GHR | Rabbit | Atlas | HPA057705 | 1:200 | IF |
| TET1 | Mouse | GeneTex | GTX627420 | 1:200 | IF |
| mCherry | Rabbit | abcam | ab167453 | 1:200 | IF |
| ALB | Mouse | R&D | MAB1455 | 1:200 | IF |
| ALB | Goat | Bethyl | A80-129A | 1µg | ELISA |
| HNF1A | Rabbit | Atlas | HPA035231 | 1:200 | IF |
| normal IgG control | Rabbit | abcam | ab37415 | 5µg | ChIP |

**Table S2:** List of TaqMan probes used for qPCR.

| Gene | Gene name | TaqMan probe (Catalog #) |
| --- | --- | --- |
| ACSS2 | Acyl-coenzyme A synthetase short-chain family member 2 | Hs01122829_m1 |
| ALB | Albumin | Hs00609411_m1 |
| FAH | Fumarylacetoacetate hydrolase | Hs00164611_m1 |
| ASL | Argininosuccinate lyase | Hs00902699_m1 |
| CPS1 | Carbamoyl phosphate synthase | Hs00157048_m1 |
| OTC | Ornithine transcarbamylase | Hs00166892_m1 |
| ALDH1A2 | Aldehyde dehydrogenase 1 family member A2 | Hs00180254_m1 |
| ALDH6A1 | Aldehyde dehydrogenase 6 family member A1 | Hs00194421_m1 |
| GLUL | Glutamine synthetase | Hs00365928_g1 |
| HIF1A | Hypoxia-inducible factor 1-alpha | Hs00153153_m1 |
| SREBF1 | Sterol regulatory element-binding protein 1 | Hs02561944_s1 |
| ARG1 | Arginase 1 | Hs00163660_m1 |
| GSTA2 | Glutathione S-transferase A2 | Hs00747232_mH |

**Table S3:**
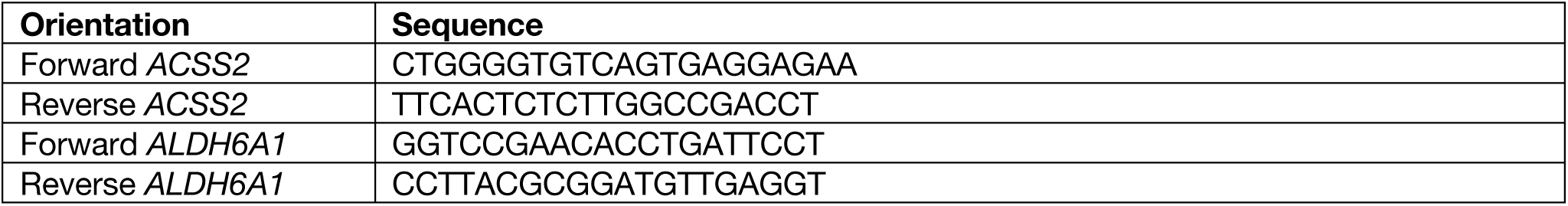
List of Custom primers used for ChIP-PCR.

**Table S4:**
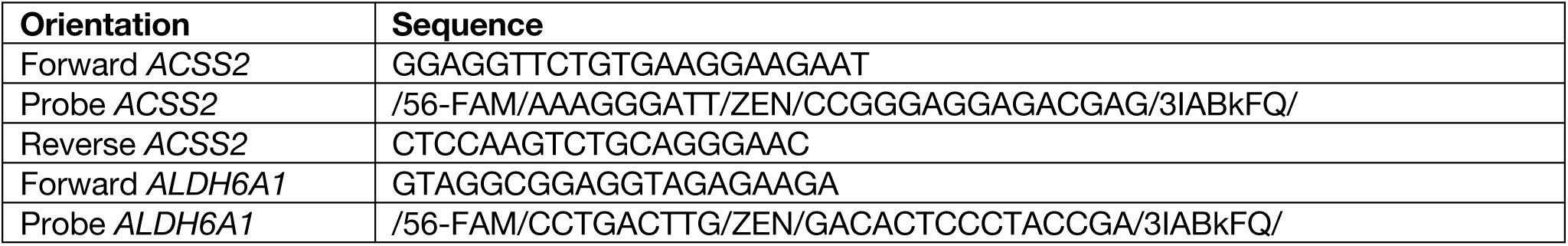
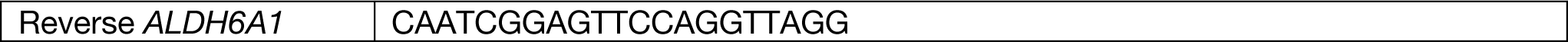
List of Custom primers used for ChIP-qPCR.

## References

1 Ben-Moshe, S. & Itzkovitz, S. Spatial heterogeneity in the mammalian liver. Nature Reviews Gastroenterology & Hepatology 16, 395–410 (2019). 10.1038/s41575-019-0134-x

2 Martini, T., Naef, F. & Tchorz, J. S. Spatiotemporal Metabolic Liver Zonation and Consequences on Pathophysiology. Annual Review of Pathology: Mechanisms of Disease 18, 439–466 (2023). 10.1146/annurev-pathmechdis-031521-024831

3 Bartl, M. et al. Optimality in the zonation of ammonia detoxification in rodent liver. Archives of Toxicology 89, 2069–2078 (2015). 10.1007/s00204-015-1596-4

4 Husson, A., Brasse-Lagnel, C., Fairand, A., Renouf, S. & Lavoinne, A. Argininosuccinate synthetase from the urea cycle to the citrulline–NO cycle. European Journal of Biochemistry 270, 1887–1899 (2003). 10.1046/j.1432-1033.2003.03559.x

5 Häussinger, D. Nitrogen metabolism in liver: structural and functional organization and physiological relevance. Biochem J 267, 281–290 (1990). 10.1042/bj2670281

6 van Straten, G. et al. Aberrant Expression and Distribution of Enzymes of the Urea Cycle and Other Ammonia Metabolizing Pathways in Dogs with Congenital Portosystemic Shunts. PLOS ONE 9, e100077 (2014). 10.1371/journal.pone.0100077

7 McNaughton, L. et al. Distribution of nitric oxide synthase in normal and cirrhotic human liver. Proceedings of the National Academy of Sciences 99, 17161–17166 (2002). doi:10.1073/pnas.0134112100

8 Baldelli, S. et al. Glutathione and Nitric Oxide: Key Team Players in Use and Disuse of Skeletal Muscle. Nutrients 11 (2019). 10.3390/nu11102318

9 Yu, Y. et al. A comparative analysis of liver transcriptome suggests divergent liver function among human, mouse and rat. Genomics 96, 281–289 (2010). 10.1016/j.ygeno.2010.08.003

10 Jiang, C. et al. Comparative Transcriptomics Analyses in Livers of Mice, Humans, and Humanized Mice Define Human-Specific Gene Networks. Cells 9, 2566 (2020).

11 Cunningham, R. P. & Porat-Shliom, N. Liver Zonation - Revisiting Old Questions With New Technologies. Front Physiol 12, 732929 (2021). 10.3389/fphys.2021.732929

12 Kolbe, E. et al. Mutual Zonated Interactions of Wnt and Hh Signaling Are Orchestrating the Metabolism of the Adult Liver in Mice and Human. Cell Rep 29, 4553–4567.e4557 (2019). 10.1016/j.celrep.2019.11.104

13 Scheidecker, B. et al. Induction of in vitro Metabolic Zonation in Primary Hepatocytes Requires Both Near-Physiological Oxygen Concentration and Flux. Frontiers in Bioengineering and Biotechnology 8 (2020).

14 McCarty, W. J., Usta, O. B. & Yarmush, M. L. A Microfabricated Platform for Generating Physiologically-Relevant Hepatocyte Zonation. Scientific Reports 6, 26868 (2016). 10.1038/srep26868

15 Mitani, S. et al. Human ESC/iPSC-Derived Hepatocyte-like Cells Achieve Zone-Specific Hepatic Properties by Modulation of WNT Signaling. Mol Ther 25, 1420–1433 (2017). 10.1016/j.ymthe.2017.04.006

16 Wahlicht, T. et al. Controlled Functional Zonation of Hepatocytes In Vitro by Engineering of Wnt Signaling. ACS Synthetic Biology 9, 1638–1649 (2020). 10.1021/acssynbio.9b00435

17 Nejak-Bowen, K. N., Zeng, G., Tan, X., Cieply, B. & Monga, S. P. Beta-catenin regulates vitamin C biosynthesis and cell survival in murine liver. J Biol Chem 284, 28115–28127 (2009). 10.1074/jbc.M109.047258

18 Bacchus, H., Heiffer, M. H. & Altszuler, N. Potentiating Effect of Ascorbic Acid on Cortisone- Induced Gluconeogenesis. Proceedings of the Society for Experimental Biology and Medicine 79, 648–650 (1952). 10.3181/00379727-79-19474

19 Greene, Y. J., Harwood, H. J. & Stacpoole, P. W. Ascorbic acid regulation of 3-hydroxy-3- methylglutaryl coenzyme A reductase activity and cholesterol synthesis in guinea pig liver. Biochimica et Biophysica Acta (BBA) - Lipids and Lipid Metabolism 834, 134–138 (1985). 10.1016/0005-2760(85)90186-9

20 Kang, J. et al. Simultaneous deletion of the methylcytosine oxidases Tet1 and Tet3 increases transcriptome variability in early embryogenesis. Proceedings of the National Academy of Sciences 112, E4236–E4245 (2015). doi:10.1073/pnas.1510510112

21 Xu, Y. et al. Ascorbate protects liver from metabolic disorder through inhibition of lipogenesis and suppressor of cytokine signaling 3 (SOCS3). Nutrition & Metabolism 17, 17 (2020). 10.1186/s12986-020-0431-y

22 Ha, T. Y., Otsuka, M. & Arakawa, N. Ascorbate Indirectly Stimulates Fatty Acid Utilization in Primary Cultured Guinea Pig Hepatocytes by Enhancing Carnitine Synthesis. The Journal of Nutrition 124, 732–737 (1994). 10.1093/jn/124.5.732

23 Creeden, J. F., Gordon, D. M., Stec, D. E. & Hinds, T. D., Jr. Bilirubin as a metabolic hormone: the physiological relevance of low levels. Am J Physiol Endocrinol Metab 320, E191–E207 (2021). 10.1152/ajpendo.00405.2020

24 Gazzin, S. et al. Bilirubin accumulation and Cyp mRNA expression in selected brain regions of jaundiced Gunn rat pups. Pediatric Research 71, 653–660 (2012). 10.1038/pr.2012.23

25 Tanii, H. et al. Induction of Cytochrome P450 2A6 by Bilirubin in Human Hepatocytes. Pharmacology & Pharmacy 04, 182–190 (2013). 10.4236/pp.2013.42026

26 Ma, R., Martínez-Ramírez, A. S., Borders, T. L., Gao, F. & Sosa-Pineda, B. Metabolic and non- metabolic liver zonation is established non-synchronously and requires sinusoidal Wnts. eLife 9, e46206 (2020). 10.7554/eLife.46206

27 Zhu, S. et al. Liver Endothelial Heg Regulates Vascular/Biliary Network Patterning and Metabolic Zonation Via Wnt Signaling. Cellular and Molecular Gastroenterology and Hepatology 13, 1757–1783 (2022). 10.1016/j.jcmgh.2022.02.010

28 Ikeda, Y. et al. Bilirubin exerts pro-angiogenic property through Akt-eNOS-dependent pathway. Hypertension Research 38, 733–740 (2015). 10.1038/hr.2015.74

29 Bandara, N. et al. Molecular control of nitric oxide synthesis through eNOS and caveolin-1 interaction regulates osteogenic differentiation of adipose-derived stem cells by modulation of Wnt/β- catenin signaling. Stem Cell Res Ther 7, 182 (2016). 10.1186/s13287-016-0442-9

30 Kim, S. G. et al. Bilirubin Activates Transcription of HIF-1α in Human Proximal Tubular Cells Cultured in the Physiologic Oxygen Content. J Korean Med Sci 29, S146–S154 (2014).

31 Cui, J., Pan, Y. H., Zhang, Y., Jones, G. & Zhang, S. Progressive pseudogenization: vitamin C synthesis and its loss in bats. Mol Biol Evol 28, 1025–1031 (2011). 10.1093/molbev/msq286

32. 32 Reza, H. A., et al. Synthetic augmentation of bilirubin metabolism in human pluripotent stem cell- derived liver organoids. Stem Cell Reports (2023). 10.1016/j.stemcr.2023.09.006

33 Horio, F., Ozaki, K., Yoshida, A., Makino, S. & Hayashi, Y. Requirement for ascorbic acid in a rat mutant unable to synthesize ascorbic acid. J Nutr 115, 1630–1640 (1985). 10.1093/jn/115.12.1630

34 Kimura, M. et al. En masse organoid phenotyping informs metabolic-associated genetic susceptibility to NASH. Cell 185, 4216–4232 e4216 (2022). 10.1016/j.cell.2022.09.031

35 Ouchi, R. et al. Modeling Steatohepatitis in Humans with Pluripotent Stem Cell-Derived Organoids. Cell Metab 30, 374–384 e376 (2019). 10.1016/j.cmet.2019.05.007

36 Shinozawa, T. et al. High-Fidelity Drug-Induced Liver Injury Screen Using Human Pluripotent Stem Cell-Derived Organoids. Gastroenterology 160, 831–846 e810 (2021). 10.1053/j.gastro.2020.10.002

37 Andrews, T. S. et al. Single-Cell, Single-Nucleus, and Spatial RNA Sequencing of the Human Liver Identifies Cholangiocyte and Mesenchymal Heterogeneity. Hepatology Communications 6, 821–840 (2022). 10.1002/hep4.1854

38 Aizarani, N. et al. A human liver cell atlas reveals heterogeneity and epithelial progenitors. Nature 572, 199–204 (2019). 10.1038/s41586-019-1373-2

39 MacParland, S. A. et al. Single cell RNA sequencing of human liver reveals distinct intrahepatic macrophage populations. Nature Communications 9, 4383 (2018). 10.1038/s41467-018-06318-7

40 Wesley, B. T. et al. Single-cell atlas of human liver development reveals pathways directing hepatic cell fates. Nature Cell Biology 24, 1487–1498 (2022). 10.1038/s41556-022-00989-7

41 Camp, J. G. et al. Multilineage communication regulates human liver bud development from pluripotency. Nature 546, 533–538 (2017).

42 Guan, Y. et al. A human multi-lineage hepatic organoid model for liver fibrosis. Nature communications 12, 6138 (2021).

43 Harrison, S. P. et al. Scalable production of tissue-like vascularized liver organoids from human PSCs. Experimental & Molecular Medicine 55, 2005–2024 (2023).

44 Hess, A. et al. Single-cell transcriptomics stratifies organoid models of metabolic dysfunction- associated steatotic liver disease. The EMBO Journal 42, e113898 (2023).

45 Zhang, C. J. et al. A human liver organoid screening platform for DILI risk prediction. Journal of hepatology 78, 998–1006 (2023).

46 Dann, E., Henderson, N. C., Teichmann, S. A., Morgan, M. D. & Marioni, J. C. Differential abundance testing on single-cell data using k-nearest neighbor graphs. Nature biotechnology 40, 245–253 (2022).

47 Ton, M.-L. N. et al. An atlas of rabbit development as a model for single-cell comparative genomics. Nature Cell Biology 25, 1061–1072 (2023).

48 Xu, C.-R. et al. Chromatin &#x201c;Prepattern&#x201d; and Histone Modifiers in a Fate Choice for Liver and Pancreas. Science 332, 963–966 (2011). doi:10.1126/science.1202845

49 Thakur, A. et al. Hepatocyte Nuclear Factor 4-Alpha Is Essential for the Active Epigenetic State at Enhancers in Mouse Liver. Hepatology 70, 1360–1376 (2019). 10.1002/hep.30631

50 Raisner, R. et al. Enhancer Activity Requires CBP/P300 Bromodomain-Dependent Histone H3K27 Acetylation. Cell Reports 24, 1722–1729 (2018). 10.1016/j.celrep.2018.07.041

51 Kapadia, B. et al. PIMT regulates Hepatic Gluconeogenesis in Mice. iScience, 106120 (2023). 10.1016/j.isci.2023.106120

52 Wang, Y., Viscarra, J., Kim, S. J. & Sul, H. S. Transcriptional regulation of hepatic lipogenesis. Nat Rev Mol Cell Biol 16, 678–689 (2015). 10.1038/nrm4074

53 Smith, R. P. et al. Genome-Wide Discovery of Drug-Dependent Human Liver Regulatory Elements. PLOS Genetics 10, e1004648 (2014). 10.1371/journal.pgen.1004648

54 Ölander, M. et al. Hepatocyte size fractionation allows dissection of human liver zonation. Journal of Cellular Physiology 236, 5885–5894 (2021). 10.1002/jcp.30273

55 Kang, S. W. S. et al. A spatial map of hepatic mitochondria uncovers functional heterogeneity shaped by nutrient-sensing signaling. Nature Communications 15, 1799 (2024). 10.1038/s41467-024-45751-9

56 Mariotti, V., Strazzabosco, M., Fabris, L. & Calvisi, D. F. Animal models of biliary injury and altered bile acid metabolism. Biochimica et Biophysica Acta (BBA) - Molecular Basis of Disease 1864, 1254–1261 (2018). 10.1016/j.bbadis.2017.06.027

57 Claeys, W. et al. A mouse model of hepatic encephalopathy: bile duct ligation induces brain ammonia overload, glial cell activation and neuroinflammation. Scientific Reports 12, 17558 (2022). 10.1038/s41598-022-22423-6

58 Paris, J. & Henderson, N. C. Liver zonation, revisited. Hepatology 76 (2022).

59 Bartl, M. et al.

60 Harrison, S. P. et al. Liver Organoids: Recent Developments, Limitations and Potential. Frontiers in Medicine 8 (2021). 10.3389/fmed.2021.574047

61 Wei, Y. et al. Liver homeostasis is maintained by midlobular zone 2 hepatocytes. Science 371, eabb1625 (2021). doi:10.1126/science.abb1625

62 Wu, H., Kirita, Y., Donnelly, E. L. & Humphreys, B. D. Advantages of Single-Nucleus over Single- Cell RNA Sequencing of Adult Kidney: Rare Cell Types and Novel Cell States Revealed in Fibrosis. J Am Soc Nephrol 30, 23–32 (2019). 10.1681/asn.2018090912

63 He, L. et al. Proliferation tracing reveals regional hepatocyte generation in liver homeostasis and repair. Science 371, eabc4346 (2021). doi:10.1126/science.abc4346

64 Li, W., Li, L. & Hui, L. Cell Plasticity in Liver Regeneration. Trends in Cell Biology 30, 329–338 (2020). 10.1016/j.tcb.2020.01.007

65 Brosch, M. et al. Epigenomic map of human liver reveals principles of zonated morphogenic and metabolic control. Nature Communications 9, 4150 (2018). 10.1038/s41467-018-06611-5

66 Kapadia, B. et al. PIMT regulates hepatic gluconeogenesis in mice. iScience 26, 106120 (2023). 10.1016/j.isci.2023.106120

67 Moffett, J. R., Puthillathu, N., Vengilote, R., Jaworski, D. M. & Namboodiri, A. M. Acetate Revisited: A Key Biomolecule at the Nexus of Metabolism, Epigenetics and Oncogenesis—Part 1: Acetyl-CoA, Acetogenesis and Acyl-CoA Short-Chain Synthetases. Frontiers in Physiology 11 (2020). 10.3389/fphys.2020.580167

68 Huang, H. et al. p300-Mediated Lysine 2-Hydroxyisobutyrylation Regulates Glycolysis. Mol Cell 70, 663–678.e666 (2018). 10.1016/j.molcel.2018.04.011

69 Gang, X. et al. P300 acetyltransferase regulates fatty acid synthase expression, lipid metabolism and prostate cancer growth. Oncotarget 7, 15135–15149 (2016). 10.18632/oncotarget.7715

70 Xu, W. et al. Hypoxia activates Wnt/β-catenin signaling by regulating the expression of BCL9 in human hepatocellular carcinoma. Scientific Reports 7, 40446 (2017). 10.1038/srep40446

71 Kietzmann, T. Metabolic zonation of the liver: The oxygen gradient revisited. Redox Biol 11, 622–630 (2017). 10.1016/j.redox.2017.01.012

72 Iansante, V., Mitry, R. R., Filippi, C., Fitzpatrick, E. & Dhawan, A. Human hepatocyte transplantation for liver disease: current status and future perspectives. Pediatric Research 83, 232–240 (2018). 10.1038/pr.2017.284

73 Reza, H. A., Okabe, R. & Takebe, T. Organoid transplant approaches for the liver. Transplant International 34, 2031–2045 (2021). 10.1111/tri.14128

74 He, L. et al. Transcriptional co-activator p300 maintains basal hepatic gluconeogenesis. J Biol Chem 287, 32069–32077 (2012). 10.1074/jbc.M112.385864

